# cytoFlagR: A comprehensive framework to objectively assess high-parameter cytometry data for batch effects

**DOI:** 10.1101/2025.05.27.656370

**Authors:** Shruti Eswar, Zachary T. Koenig, Amanda R. Tursi, José Cobeña-Reyes, Tamara Tilburgs, Sandra Andorf

**Author notes:** **Corresponding Author:** Sandra Andorf, PhD, Division of Biomedical Informatics, Cincinnati Children’s Hospital Medical Center University of Cincinnati, 3333 Burnet Avenue, Cincinnati, OH 45229. Shruti Eswar and Zachary T. Koenig contributed equally as first authors. Tamara Tilburgs and Sandra Andorf contributed equally as senior authors.

## Abstract

**Motivation:** High-parameter cytometry is widely used in longitudinal studies, but technical variation across batches can confound biological signals. However, tools that objectively identify problematic batches and markers are limited.

**Results:** We introduce cytoFlagR, a comprehensive tool to flag batch-related problems at the marker and cell cluster level based on robust statistical evaluations. Batch and marker variations are assessed based on median signal intensities of negative and positive cell populations and positive cell frequencies, along with Earth Mover’s Distance (EMD) of signal intensity distributions. Additionally, cytoFlagR identifies cell type specific batch problems via unsupervised clustering and is suitable for mass and spectral cytometry datasets where it objectively detects distinct types of batch issues. We demonstrated cytoFlagR’s utility for assessing datasets that include or lack reference controls. Thus, cytoFlagR improves quality control by objective identification of technical variations that may impact downstream analysis.

**Availability and Implementation:** cytoFlagR is freely available as R scripts with documentation and an example at https://github.com/AndorfLab/cytoFlagR.

## 1 Introduction

High-parameter cytometry techniques are increasingly used in large, longitudinal immunological and clinical studies to enable the study of biological mechanisms and immune regulations. Over the last decades, advancement in technologies such as mass cytometry (CyTOF) and spectral flow cytometry have enabled the measurement of over 40 cellular markers for millions of single cells per sample (Spitzer and Nolan 2016; McKinnon 2018). However, collecting cytometry data over a long period of time can result in technical variations between the acquisition batches, driven by several factors including differences in sample preparation, instrument-dependent factors, reagents used and other technical artifacts (Leipold, Newell and Maecker 2015; Van Gassen *et al*. 2020a; Cossarizza *et al*. 2021). This technical variability between acquisition runs, osen referred to as batch effects, can confound biological signals in the experimental data (Lo *et al*. 2022). Batch effects in cytometry can impact individual markers differently, and sometimes technical variations between batches are not detectable on a global scale but only within specific cell populations (Van Gassen *et al*. 2020a).

To minimize the impact of batch effects, several different approaches exist to adjust the data before downstream analysis, either by utilizing reference control samples that were included in each batch (Schuyler *et al*. 2019; Trussart *et al*. 2020; Van Gassen *et al*. 2020b; Pedersen *et al*. 2022), or through approaches that do not rely on such controls (Hahne *et al*. 2010; Ogishi *et al*. 2021; Pedersen *et al*. 2022; Quintelier *et al*. 2025). However, there is currently a lack of robust statistical approaches that objectively assess whether and to what extent a cytometry dataset is affected by technical variations.

Here, we present cytoFlagR, a comprehensive, objective, and interpretable approach to assess batch effects in cytometry datasets. cytoFlagR uses a set of robust statistical metrics to identify markers and batches that are significant outliers and potentially problematic at the global marker and/or cell population level. It assesses shiss in medians of marker intensity distributions of the positive and negative cell populations, differences in cell frequencies of the positive population, and Earth Mover’s Distance (EMD)-based comparisons of signal intensity distributions. Additionally, cytoFlagR performs unsupervised clustering of the control samples to evaluate whether problematic batches impact the detection of canonical cell populations. This tool was developed to utilize reference control samples, but we demonstrated its utility in assessing a mass cytometry dataset for which no control samples were included. In summary, cytoFlagR enables users to identify distinct types of technical variations that may confound downstream analyses. By identifying these variations, the tool supports users in their decision regarding applying batch correction or excluding batches or markers from subsequent analyses.

## 2 Methods

### 2.1. cytoFlagR workflow

The tool relies only on the control samples that were included in each experimental batch (Fig. 1, box A). The input for cytoFlagR consists of pre-gated live, single cells from each of the control samples across all batches which are then transformed using an inverse hyperbolic sine (arcsinh) function. While cytoFlagR primarily utilizes control samples, it can also be applied to biological samples, provided the user accounts for the inherent biological variability between samples when interpreting the tool’s outputs. In addition to common visual assessments (Fig. 1, box B, details in Supplementary methods), cytoFlagR uses two main methods on a per-marker basis for every control to flag possibly problematic batches and markers: an Inter-Quartile Range (IQR) based assessment and an Earth Mover’s Distance (EMD) based assessment.

**Figure 1.**
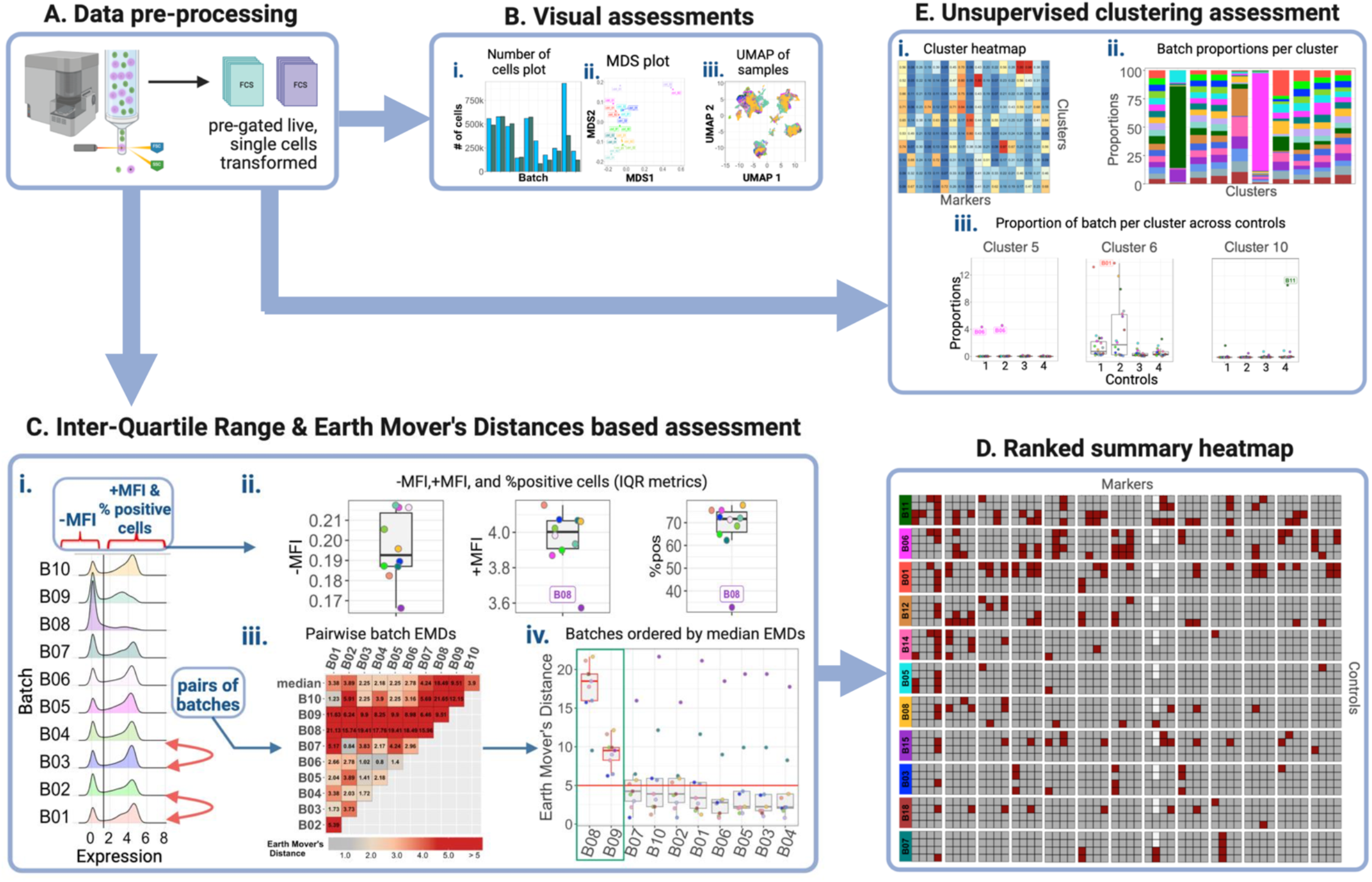
Overview of cytoFlagR. The workflow consists of five main steps, typically performed on control samples included in each batch of the experiment: **(A)** Transformation of FCS files containing pre-gated live single cells. **(B)** Initial visual inspection of batch effects is carried out using: (i) a bar graph showing the number of cells in each batch for every control sample, (ii) a Multi-Dimensional Scaling (MDS) plot, and (iii) a Uniform Manifold Approximation and Projection (UMAP) of the controls colored by the batches. **(C)** For every marker and control, potential batch issues are detected using four metrics derived from two main approaches: an Inter-Quartile Range (IQR) based assessment and an Earth Mover’s Distance (EMD) based metric. (i) The density distributions of one marker for every control across all batches are visualized and a threshold (black line) is determined to divide the distributions into negative and positive populations. (ii) An IQR-based assessment identifies potentially problematic batches based on negative and positive median fluorescence intensities (MFIs) and the percent of positive cells (%pos). All metric values are visualized in boxplots with flagged batches labeled. (iii) The EMD-based metric assesses the shape of full distributions. Pairwise EMDs are calculated between all batches per control per marker and visualized as a heatmap along with the median EMD per batch. (iv) Boxplots of these pairwise EMDs ordered by their medians are used to flag problematic batches. **(D)** A summarized heatmap integrates results from all four metrics across all markers and control samples. Markers and batches are ranked based on the frequency of their flags, from highest number of flags to lowest. **(E)** To evaluate the impact of potential problematic batches on the overall population structure, unsupervised clustering (FlowSOM) is performed on all control samples. (i) A cluster heatmap of marker expressions. (ii) The proportion of analyzed cells of each batch present in a cluster is calculated and visualized. (iii) For clusters containing potentially problematic batches, cell proportions across individual controls are visualized and assessed. Created in BioRender. E, S. (2025) https://BioRender.com/8dh5nmv

We use the term *control sample* or *control* throughout this manuscript to refer to one of the standardized reference samples included in each experimental batch. It is assumed that each control was included only once per batch. Therefore, when referring to a specific instance of a control included in a particular batch, we will refer to it by the corresponding batch number.

#### 2.1.1. Inter-Quartile Range (IQR)-based assessment

Using the density distributions of expression values across batches for each marker and control, cytoFlagR automatically determines a threshold value (see Supplementary methods) to divide the data into negative and positive populations (Fig. 1, box C (i)). The user can adjust the automatically determined threshold.

The negative and positive populations are used to calculate the three IQR-based metrics used by cytoFlagR: 1. Negative median fluorescence (or signal in case of mass cytometry) intensities (-MFI), 2. positive median fluorescence intensities (+MFI), 3. percent of cells in the positive population (%pos). These metrics are calculated and assessed per control sample per marker across all batches. For each metric, a batch is flagged as potentially showing a batch effect if the metric value for the corresponding control sample is more than 1.5 IQR above the 3rd quartile or below the 1st quartile, following the outlier definition by (McGill, W.Tukey and Larsen 1978). To ensure reliability of batches flagged by the MFI metrics, any instance with less than 100 cells present in the negative or positive population is excluded from the respective IQR-based assessment. Boxplots of the three metrics (-MFI, +MFI, and %pos) are generated for each marker and control, with batches that were determined as problematic highlighted. The plots are generated with the y-axis being variable per marker (Fig. 1, box C (ii)) and with the y-axis fixed to the same range across all markers.

#### 2.1.3. Earth Mover’s Distance (EMD)-based assessment

To identify dissimilarities in the distribution of expression values across batches, the Earth Mover’s Distance (EMD) (Rubner, Tomasi and Guibas 2000) is calculated for each pair of batches, separately for each marker and control sample (Fig. 1, box C (i)). To generate the fourth assessment metric per batch, the median EMD of each batch, i.e. the median of its pairwise distances to all other batches for that marker and control, is computed. This batch-level median EMD metric is then used to flag potentially problematic batches (default threshold of ≥5). Additional details are provided in the Supplementary methods.

The pairwise distances are visualized as heatmaps (Fig. 1, box C (iii)), with the top row representing the median of the pairwise EMDs for that batch. Additionally, boxplots of pairwise EMDs ordered by their median EMD are generated (Fig. 1, box C (iv)), with flagged batches highlighted in red.

#### 2.1.4. Ranked summary of flags

A heatmap summarizes the batches and markers flagged as potentially exhibiting batch effects in the controls, based on the four metrics (-MFI, +MFI, %pos, and median EMD) (Fig. 1, box D). The batches and markers are ordered from the largest to the lowest number of flags across all controls.

#### 2.1.5. Unsupervised clustering-based assessment

To assess whether any batches exhibit a batch effect that might impact downstream cell population quantification, cytoFlagR includes an additional assessment based on unsupervised clustering. This assessment relies on the expectation that in a dataset without batch effects, cells from the control sample included in different batches will be evenly distributed within each cluster (i.e., cell population). For example, in a dataset with 10 batches, each cluster should contain about 10% of its cells from each batch, assuming each batch contributes the same number of total cells.

In this assessment, control samples from all batches (randomly down-sampled to 20,000 cells per file) are clustered using FlowSOM (R package *FlowSOM, v2*.*12*.*0*) (Van Gassen *et al*. 2015). An appropriate meta-cluster number to represent canonical cell populations is set by the user (R package *ConsensusClusterPlus v1*.*68*.*0*) (Wilkerson and Hayes 2010). The clusters are visualized using marker expression heatmaps (Fig. 1, box E (i)).

Proportions of the number of cells per batch in each cluster are visualized in stacked bar graphs (Fig. 1, box E (ii)). If the contribution of cells from one or more batches in one cluster is greater than 2 times the expected proportion based on the total number of batches, these batches are considered to exhibit a batch effect. Next, these clusters are further assessed to determine if a specific control sample drives the potential batch effect, utilizing boxplots showing the per-batch contributions stratified by control. A batch is flagged as potentially problematic if its contribution exceeds 1.5 times the expected proportion in that cluster for the control (Fig. 1, box E (iii)).

### 2.2. Evaluations

To evaluate the robustness and usability of cytoFlagR, several analyses were performed to assess: i. the impact of the number of batches in a dataset on cytoFlagR’s performance; ii. the impact of variable thresholds to separate the negative and positive populations, and the performance of the automated approach to determine the threshold; iii. the effect of the number of down-sampled cells on the stability of EMD calculations. See Supplementary methods for details.

### 2.3. Datasets

This tool was developed and tested on mass and spectral flow cytometry datasets (Table 1). Descriptions of the two mass cytometry datasets (Neeland *et al*. 2020; Van Gassen *et al*. 2020b; Tursi *et al*. 2022) and the original spectral cytometry dataset (Sup. Table 1) are available in the Supplementary methods.

**Table 1.**
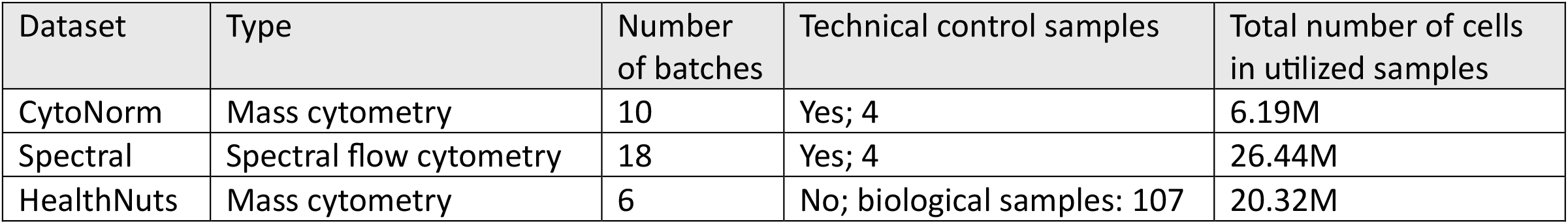
Overview of u1lized datasets. Details are provided in the supplementary methods.

| Dataset | Type | Number of batches | Technical control samples | Total number of cells in utilized samples |
| --- | --- | --- | --- | --- |
| CytoNorm | Mass cytometry | 10 | Yes; 4 | 6.19M |
| Spectral | Spectral flow cytometry | 18 | Yes; 4 | 26.44M |
| HealthNuts | Mass cytometry | 6 | No; biological samples: 107 | 20.32M |

## 3 Results

### 3.1. Illustrating cytoFlagR on commonly observed batch effect cases

The cytoFlagR tool detects distinct types of common batch related problems (batch effects) in cytometry datasets (Fig. 1). An example dataset from the HealthNuts study, consisting of 15 PBMC samples, each artificially assigned as a batch, is used for illustration. The dataset shows batch effects for the example marker in four samples, where MFIs of negative populations align across batches, but positive population MFIs or percent cells appear decreased (B11, B12, B13) or increased (B03), suggesting batch effects are present (Fig. 2A). Using cytoFlagR’s objective computational assessment based on the MFI of the negative and positive cell populations, and the percentage of positive cells, these four batches are all flagged as potentially problematic in one or more of the IQR metrics (Fig. 2B-D). The +MFI metric highlights batches B03, B11, and B12 (Fig. 2C), while the %pos metric flags B11 and B13 (Fig. 2D).

**Figure 2.**
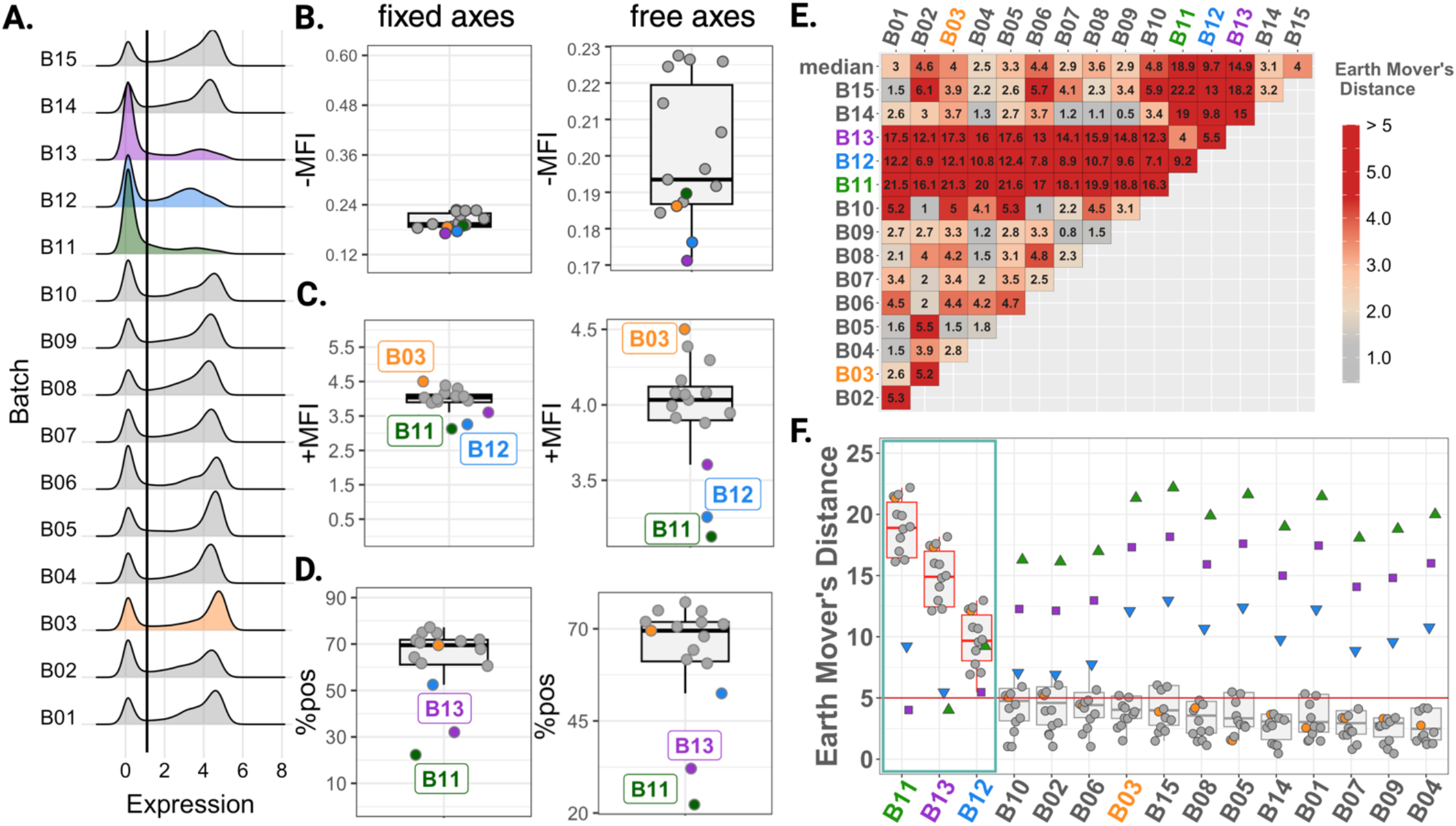
Batch effects frequently observed in cytometry datasets are detected by cytoFlagR. **(A)** Density distributions across 15 batches for one marker with a threshold value (black line) separating negative from positive populations. Batches B03, B11, B12 and B13 exhibit different batch effects. **(B-D)** Outlier batch detection results for –MFI, +MFI, and % positive cells (%pos), represented by boxplots. Y-axes ranges are fixed (left) and variable per maker (right). Batches B03, B11 and B12 are flagged as having potential batch effects in the +MFI assessment **(C)**, while batches B11 and B13, which have diminished positive peaks, are detected by the %pos outlier assessment **(D). (E)** Pairwise Earth Mover’s Distances (EMDs) across all batches, visualized as a heatmap. The top row shows the median EMD of each batch, with batches B11, B13 and B12 having a median EMD ≥ 5, flagging them as potential sources of batch effects. **(F)** Pairwise EMDs per batch, ordered by median value, are represented as boxplots. The three flagged batches are outlined in red and highlighted by a teal box. [MFI: median fluorescence intensity]

To further refine detection of batch effects in overall shapes of marker expression distributions across all batches, pairwise EMDs and median EMDs of each batch are also calculated and visualized in a heatmap (Fig. 2E) along with boxplots of pairwise EMDs ordered by the batch-level median EMD (Fig. 2F). Here batches with a low EMD are more similar to each other, while high EMDs indicate larger differences in marker expression distributions between the control samples of two batches. The heatmap and boxplots accurately flag batches B11, B12, and B13 as their median EMDs of 18.9, 9.7, and 14.9 respectively, exceed the threshold EMD of 5 used by cytoFlagR to identify potential batch effects. Thus, the combination of IQR and EMD-based flags generated by cytoFlagR accurately highlight distinct types of potential batch related technical problems.

### 3.2. Marker-wise detection of potentially problematic batches and markers within the spectral dataset

To explain the detailed workflow of cytoFlagR on a more complex dataset, we applied the tool to the spectral dataset. First, cytoFlagR creates visualizations for initial, exploratory assessment: i. The number of live, single cells per sample across each of the 18 batches, showing the variation in cell number per sample with some samples containing as lows as 23,585 live single cells (Sup. Fig. 1A); ii. The Multi-Dimensional Scaling (MDS) plot depicts clear separation between the unstimulated and stimulated samples, as expected, but also reveals additional separation of certain samples and batches, suggesting potential technical variation (Sup. Fig. 1B). Specifically, several batches (B08, B11, B14, B15, B17, B18) for control_2 were distinct from the rest in the first dimension; iii. A Uniform Manifold Approximation and Projection (UMAP) annotated with batch colors was generated to assess the uniform distribution of cells of each batch (Sup. Fig. 1C). Batch B17 exhibited clear separation from the remaining batches which is highlighted in a UMAP including density contour lines for all cells with only the cells from batch B17 depicted as dots (Sup. Fig. 1D).

Next, cytoFlagR performs detailed assessments of all batches and markers to flag samples with potential marker or batch related problems. For this, user-defined thresholds to separate the fluorescence intensity distributions into positive and negative populations for each phenotypic marker were determined. For example, the threshold for Cluster of Differentiation 3 (CD3) was set at 0.6 by the immunologists responsible for generating this dataset and applied across all four controls run in 18 batches (Fig. 3A). Then, cytoFlagR generated boxplots for the three IQR-based metrics and depicted the EMDs as heatmaps and boxplots for each control (Fig. 3B-F). CytoFlagR identified B01, B03, B06, B11 and B12 several times in the IQR and EMD metric plots, suggesting these batches are outliers and may have consistent batch related problems across different parameters. For instance, the +MFI metric (Fig. 3C) accurately highlighted batch B06 in the stimulated controls (control_3 and control_4), which exhibited a noticeable positive peak shis in the fluorescence intensity distributions (Fig. 3A). Additionally, batch B01 in the unstimulated controls (control_1 and control_2) was accurately flagged by the -MFI (Fig. 3B) and %pos metric (Fig. 3D). Samples of the unstimulated controls for that batch showed diminished positive peaks (Fig. 3A). The observable differences in the shape of the density distributions of B01, B06, B11, and B12 were reflected by the large pairwise EMDs (Fig. 3E) of these batches and elevated median EMDs (≥ 5) (Fig. 3F).

**Figure 3.**
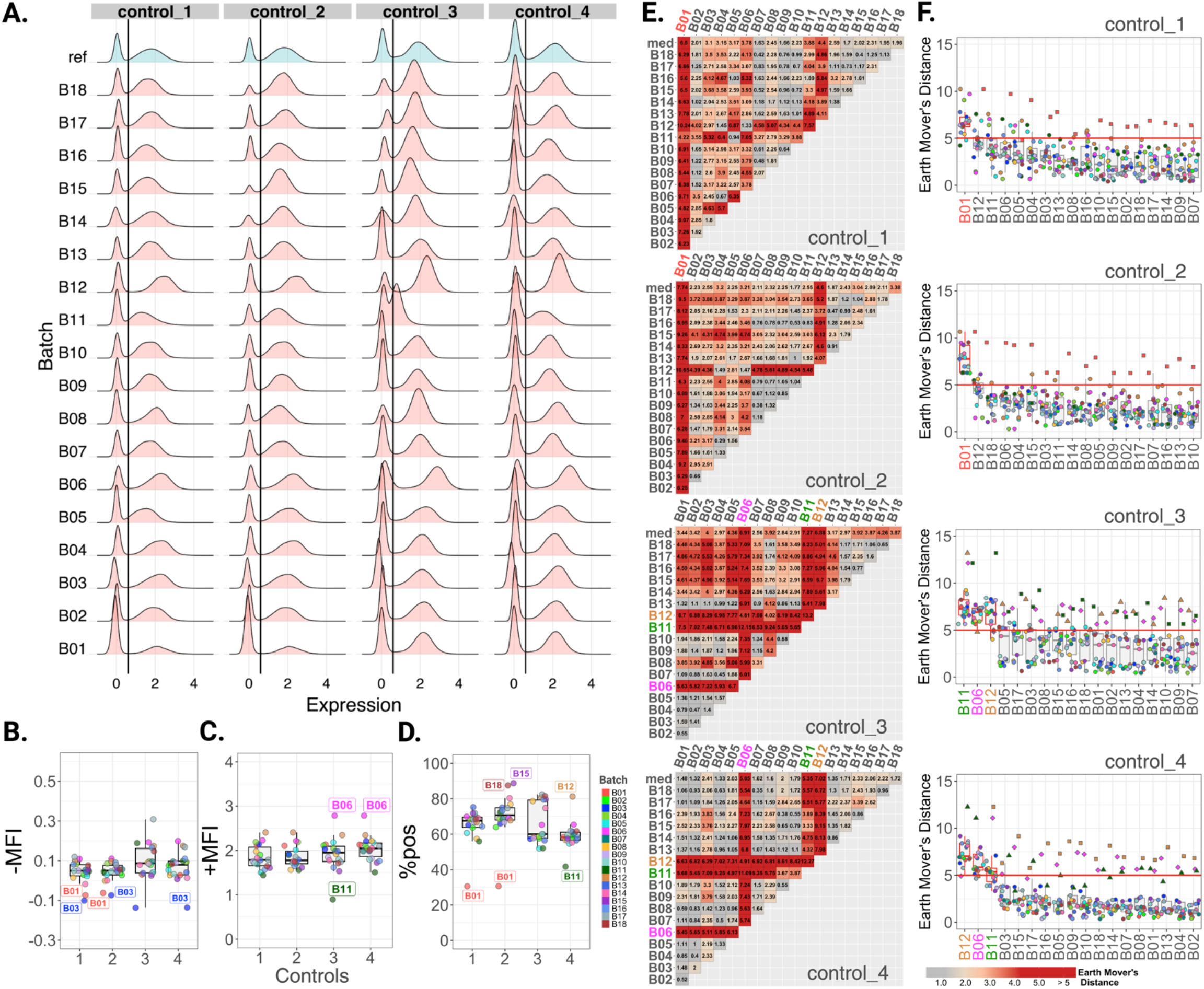
cytoFlagR highlights potenHal batch effects in the spectral cytometry dataset including four control samples across 18 batches. **(A)** Density distributions for CD3 across 18 batches of two unstimulated (control_1, control_2) and two stimulated (control_3, control_4) control samples. The black line indicates the threshold separating negative and positive populations. **(B)** Batches B01 and B03 are flagged as having potential batch effects in the unstimulated controls, while B03 is flagged in control_4 in the -MFI assessment. **(C)** The +MFI results suggest B06 as having a potential batch effect for the stimulated controls and B11 in control_3. **(D)** The % positive cells (%pos) metric shows B01 as a potential problematic batch in the unstimulated controls along with B15 and B18 in control_2 as well as B11 and B12 in control_4. **(E)** Pairwise and median EMDs (med) across all batches for each control are visualized as heatmaps. **(F)** Boxplots of EMDs ordered by their respective median values highlight B01 as a flagged batch for the unstimulated controls, and B06, B11, and B12 for the stimulated controls (represented by boxplots with a red outline). The red horizontal line at 5 represents the threshold for identifying batches with potential batch effects based on median EMD.

CytoFlagR was then applied to Granulysin (GNLY) to showcase results for a marker with unimodal fluorescence intensity distributions (Sup. Fig. 2A). Batch B17 was consistently flagged across the IQR-based (Sup. Fig. 2B) and EMD metrics (Sup. Fig. 2 C,D). These flags were consistent with dot plots, visualizing that GNLY expression of B17 was highly divergent from all other batches and suggesting a severe batch related problem in B17 GNLY expression. Similarly, B12 was consistently detected by the +MFI, %pos and EMD metrics in the stimulated controls, correctly highlighting the observable different positive populations of control samples in this batch (Sup. Fig. 2A,C,D). This demonstrates that cytoFlagR can detect problematic batches in both unimodal and bimodal marker expression distributions.

### 3.4. Summary heatmap illustrating IQR and EMD-based assessment metrics for the spectral dataset

The examples shown in Fig. 3 and Sup. Fig. 2 demonstrated cytoFlagR’s ability to detect potential batch effects on a per-marker basis for each sample (in this case, controls). Next, all flags identified across all markers, controls, and batches in this dataset were summarized in a heatmap and ranked from most to least flagged markers and batches across controls for all four metrics. A total of 377 flags out of 6,048 assessments were identified and distributed as follows: -MFI: 98, +MFI: 76, %pos: 110, and EMD: 93 (Fig. 4). Batches B11, B06 and B01 ranked highest by number of markers flagged. Granzyme B (GZMB) was the marker with the highest number of flags across all controls and batches. In this summary heatmap, GNLY in batch B17 clearly stood out as flagged for potential batch issues in all metrics and controls, consistent with findings in Sup. Fig. 2. These findings highlight cytoFlagR’s ability to provide users with a clear and comprehensive overview of potential marker- and batch-specific issues across large datasets.

**Figure 4.**
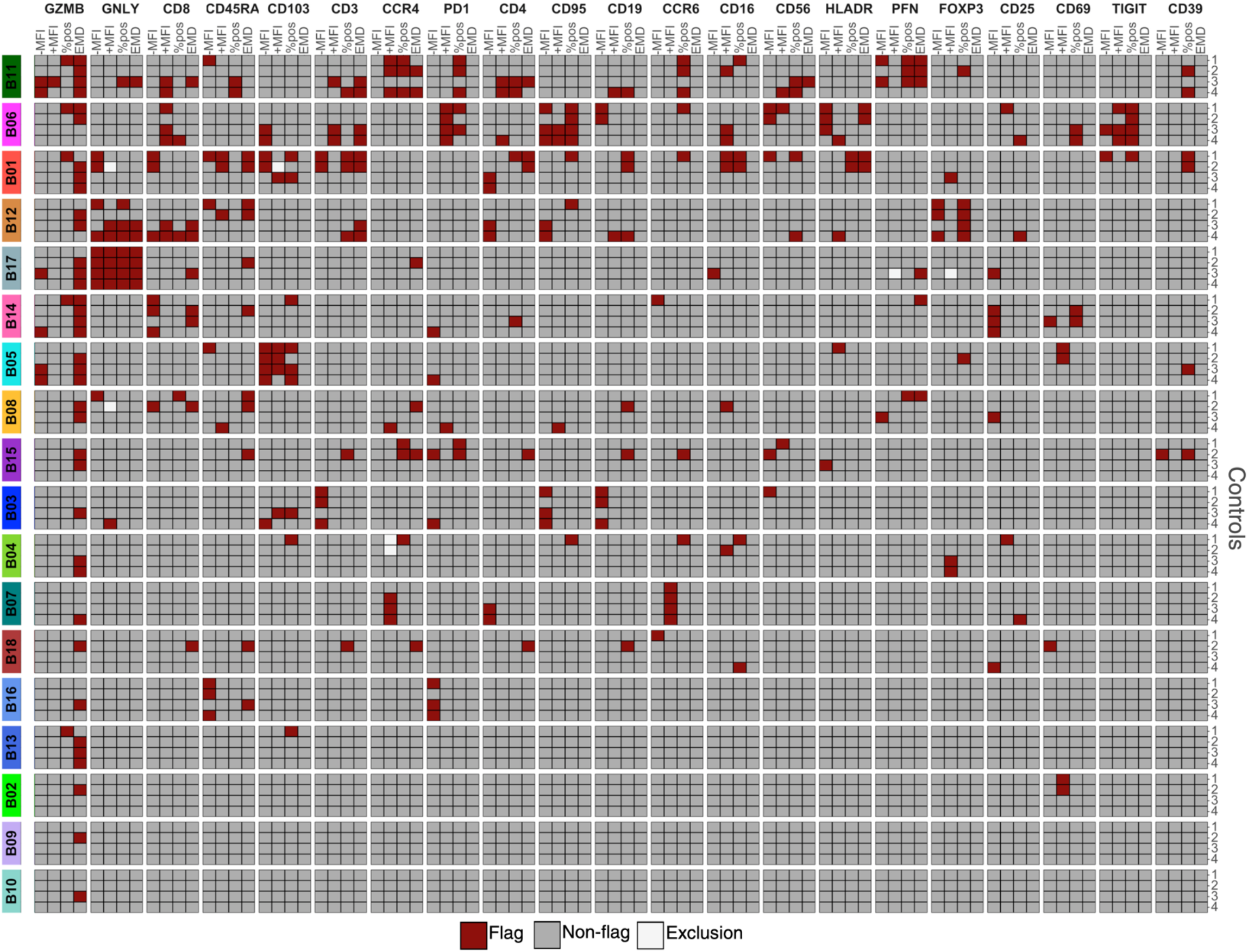
Heatmap summarizing potenHal batch effects flagged across markers, controls, and batches in the spectral cytometry dataset. The spectral cytometry dataset consisting of 4 controls, 18 batches and 21 protein channels was assessed using cytoFlagR to identify potential batch effects across four metrics (-MFI, +MFI, %pos, and EMD). Potential batch effects identified for each marker, control, and batch are summarized and visualized with a heatmap, ranking batches and markers from most frequently flagged to least. Dark red tiles indicate flagged potential batch effects, gray tiles denote no detected batch effects, and white tiles represent controls excluded from the +MFI metric assessment (fewer than 100 cells, see Methods). Batches B11, B06, and B01 show the most potential batch effects across markers and controls.

### 3.5. Assessing the impact of problematic batches on clustering of the spectral dataset

To highlight batch effects that impact downstream clustering of the data into distinct cell populations, which may not have been detected from evaluating single markers independently, cytoFlagR also identified potentially problematic batches and samples based on unsupervised clustering across all batches, control samples and selected markers. The control samples from the spectral dataset were clustered into 10 clusters using FlowSOM, represented in a heatmap of scaled median expressions, and the proportion of cells from each batch contributing to each cluster were calculated and depicted (Fig. 5A). Batches B01, B11, B06 and B12 contributed significantly more than two times the expected proportion of cells in some clusters (B01 in Cluster 3; B11 in Cluster 4; B06 in Cluster 8; and B12 in Cluster 10) (Fig. 5A,B). Analysis of the cell proportions per batch, stratified by control, highlighted which of the controls showed the batch effects in these clusters (Fig. 5B). In contrast, other clusters had a nearly equal proportion of cells from each batch (Fig 5B., clusters 1, 2, and 9, and Sup. Fig. 3, clusters 5, 6, 7). A UMAP projection of the clustered data, colored by batches, also demonstrated noticeably different and un-even batch mixing in Clusters 3, 4, 8 and 10, which supported the outcomes of this assessment (Fig. 5C). Thus, batches B01, B11, B06 and B12 which were ranked as the highest flagged batches in Figure 4 were also identified as potentially problematic batches that drive unique clusters obtained using FlowSOM.

**Figure 5.**
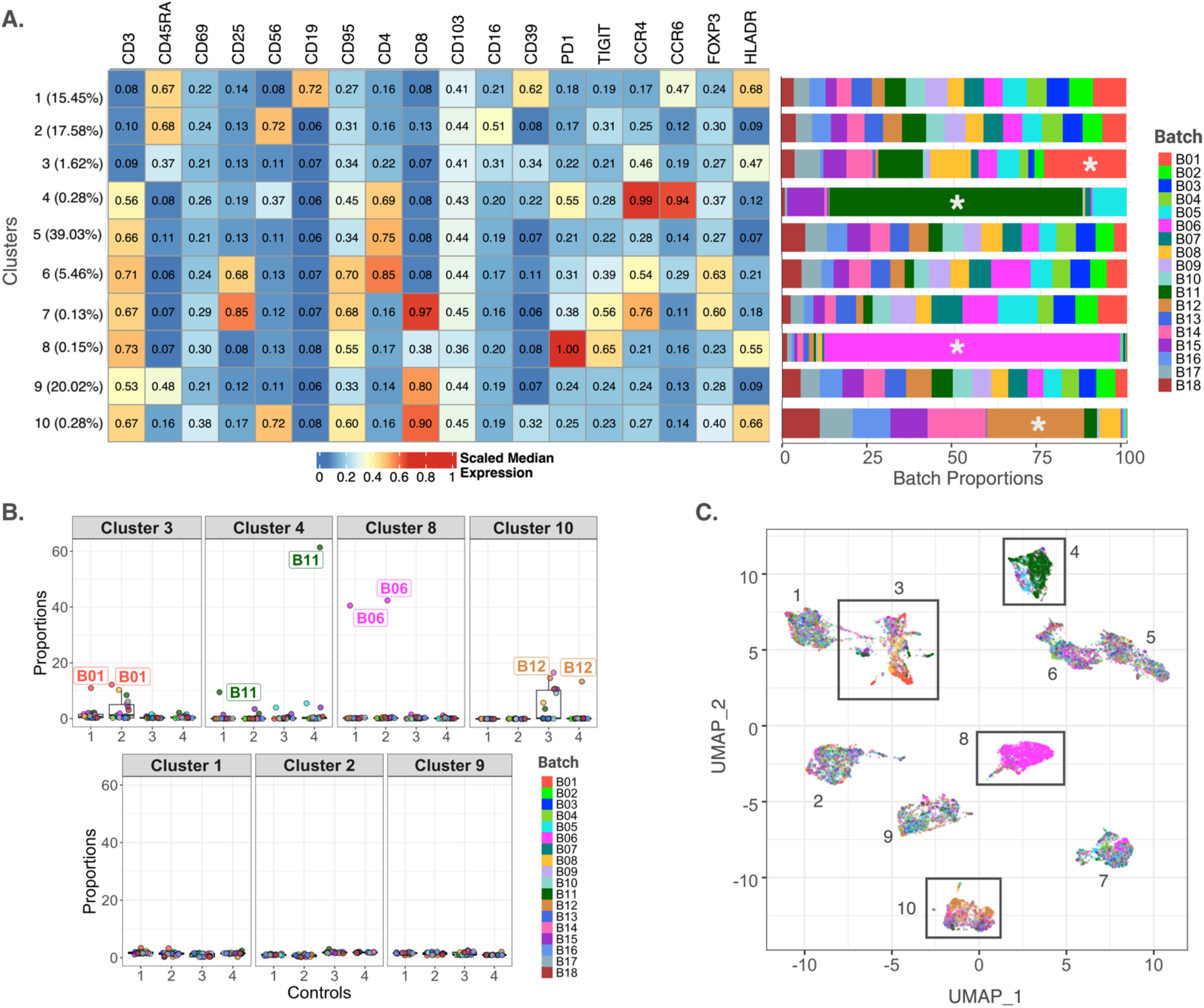
Flagging problemaHc batches based on unsupervised clustering of the spectral dataset. **(A)** Heatmap of scaled median marker expressions across 18 markers and 10 clusters (left) and the batch proportions of analyzed cells in each cluster represented by the bar plot (right). Clusters 3, 4, 8 and 10 have batches with higher proportion of cells than expected (highlighted by *) and the remaining clusters have even distribution of cells per batch. **(B)** Clusters with higher-than-expected batch proportions (see methods for definition) are visualized (top row) as boxplots of proportions stratified by controls. Cluster 3 indicates a higher abundance of batch B01 in the unstimulated controls (1 and 2). Similarly, B11 in controls 1 and 4 of Cluster 4, B06 in the unstimulated controls of Cluster 8, and batch B12 in the stimulated controls of Cluster 10 shows higher abundances in the respective clusters. Clusters 1, 2 and 9 (bottom row) are examples of clusters without problematic batches present. They have a uniform distribution of batch proportions. **(C)** Uniform Manifold Approximation and Projection (UMAP) of clusters overlayed with batch colors are consistent with the results in (A) and (B).

### 3.6. Inspection of the CytoNorm mass cytometry dataset using cytoFlagR

To show the effectiveness of cytoFlagR for datasets from a different cytometry technique, we analyzed the CytoNorm dataset, which was generated by mass cytometry (Table 1). CD25 (Sup. Fig. 4) and CD66 (Sup. Fig. 5) were assessed by cytoFlagR and detailed results are presented in the Supplementary results. In short, for CD25, two of the batches were consistently flagged as potentially problematic for control_2 across the three IQR-based and the EMD metric, reflecting the observed differences in the marker expression distributions. CD66 represents an example in which 6 batches showed very distinct marker expression distributions from the 4 remaining batches, across all 4 controls. This resulted in a wide spread of values for the three IQR-based metrics and none of the batches being flagged as potentially problematic (Sup. Fig. 5B). However, such cases of a dataset containing two sets of batches that show marker expression distributions that are distinct from each other but consistent within one set, are nevertheless detected by the EMD-based assessment. In this case, the 4 batches from the smaller set are consistently flagged as exhibiting potential batch effects across all 4 controls in the EMD assessment (Sup. Fig. 5C,D). This is also reflected in the summary of the flagged batches and markers across the 4 metrics (Sup. Fig. 6). Interestingly, the assessment based on unsupervised clustering of the control samples highlighted an additional batch as potentially problematic, mainly driven by control 4 (Sup. Fig. 7). While developed on spectral cytometry data, these results show that cytoFlagR can detect technical problems in data generated by mass cytometry.

### 3.7. Assessing biological samples of a mass cytometry dataset using cytoFlagR

To demonstrate that cytoFlagR can also be used to assess cytometry datasets which lack control samples for technical concerns, the tool was next applied to biological samples from the HealthNuts dataset, using CD27 as an example marker. Marker expression densities of the biological samples showed very similar shapes across samples within a batch, but samples within batches B03, B04, and B05 had overall different distributions from samples in other batches (Sup. Fig. 8). To account for biological variability between samples in a batch, a pooled sample per batch per stimulation condition was generated by randomly selecting and concatenating 45,000 cells per corresponding file. Marker expression densities of the pooled samples reflected the distribution shapes as seen for the individual files per batch (Sup. Fig. 9A). Given the small number of batches and the clear separation of batches into two sets, the IQR-based assessments were not successful in detecting most of the problematic batches (Sup. Fig. 9B). However, the EMD metric flagged batches B04 and B05 as potentially problematic for two of the stimulations (Sup. Fig. 9C, D). For the PMA stimulated samples, exactly half of the batches showed a distinct marker expression distribution from the other half, resulting in all batches being flagged by the median EMDs (Sup. Fig. 9D). However, the heatmap of pairwise EMDs depicted the presence of two sets of batches that each showed a small EMD between batches within a set, but a large EMD value to batches from the other set (Sup. Fig. 9C). Assessing the biological samples individually highlighted similar markers and batches as most flagged as when evaluating the dataset aser pooling samples within one batch per stimulation (Sup. Fig. 10). These results demonstrate that cytoFlagR is also able to detect problematic batches in datasets that only contain biological samples. However, while interpretating the findings, biological differences must be considered, especially when assessing individual rather than pooled samples.

### 3.8. Quantitative analysis of the effects of number of batches on marker-wise assessment metrics

The impact of the number of batches on the performance of each assessment metric used by cytoFlagR was investigated. In the spectral dataset, the detection rate of flagged cases decreased across all metrics as the number of batches was reduced from all 18 batches (i.e. all detected flags, 100% detection rate) to 2 batches (Fig. 6). The cases flagged as probable batch effects by the IQR-based metrics (-MFI, +MFI, %pos) exhibited a steady decline in the detection rate, ultimately leading to a significant drop when less than 5 batches were included and reaching 0% for 3 batches. In contrast, the EMD metric demonstrated a more gradual reduction in the detection rate, still identifying on average 74.49% of the originally flagged cases when only 2 batches were present. The CytoNorm dataset showed a similar pattern (Sup. Fig. 11). These results highlight the greater robustness of the EMD-based metric when having a small number of batches, compared to the IQR-based metrics, which are more sensitive to the overall number of batches available for comparison.

**Figure 6.**
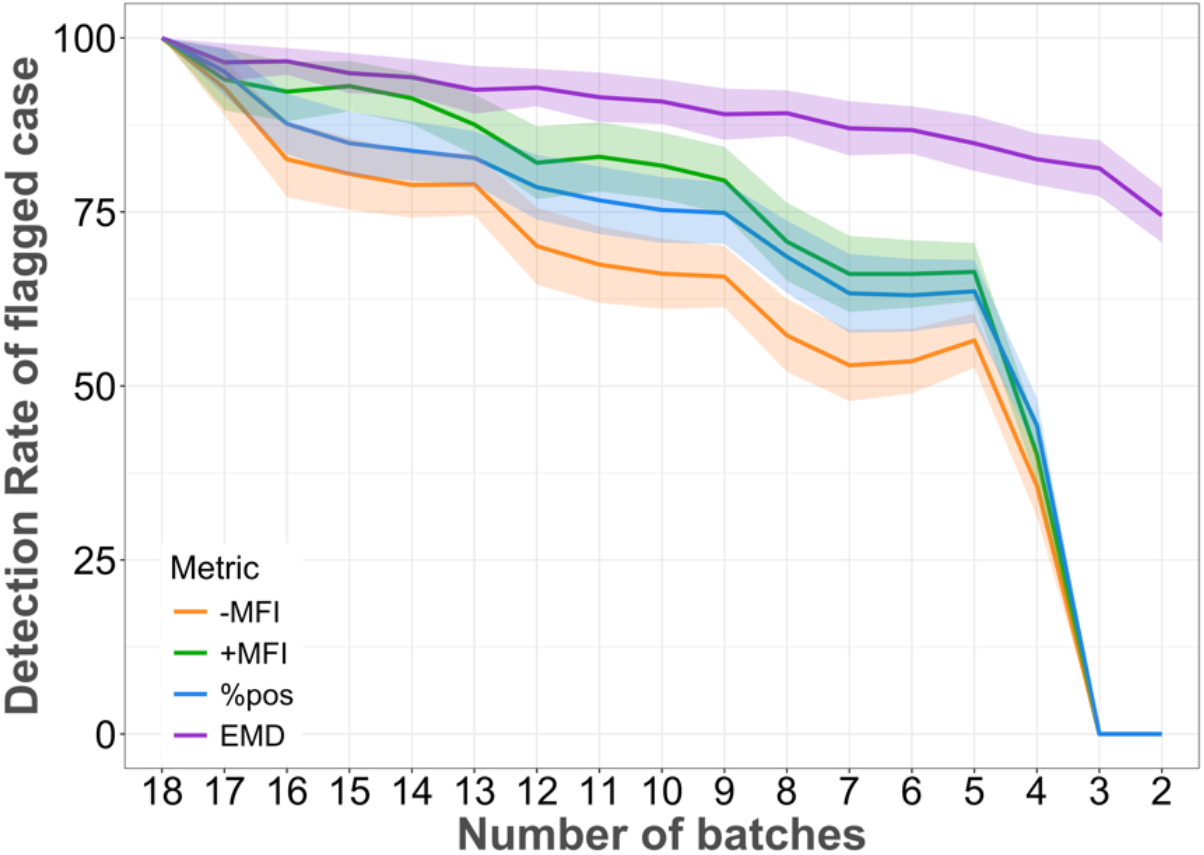
Influence of number of batches used for each assessment metric used by cytoFlagR. The lines represent the mean detection rate of flagged cases using different number of batches for each of the four metrics [-MFI, n = 89 (total cases flagged for the complete dataset at 18 batches); +MFI, n = 63; %pos, n = 103; EMD, n = 81]. 95% confidence intervals based on 50 iterations of sequentially reducing the number of batches in random order are shown as shaded areas. The cases flagged by -MFI, +MFI, and %pos metrics exhibit a downward trend, culminating in a sharp decline to zero. In contrast, the EMD metric has a more gradually decreasing trend, thereby highlighting the impact the number of batches has on the assessment metric.

### 3.9. Evaluating the impact of inter-expert variability of marker thresholds on IQR-based metrics and comparison with automated marker threshold determination

Thresholds determined to separate cytometry data into positive and negative populations for a given marker exhibit a degree of subjectivity and variability among experts (Liu *et al*. 2025). Therefore, we assessed the impact of the variability in setting the threshold across experts on the IQR-based metric values (-MFI, +MFI and %pos), as well as the difference of these values based on consensus expert thresholds when compared with cytoFlagR’s calculated automatically thresholds. The IQR-based metric values showed strong correlations when using the lowest and highest expert defined thresholds, with slightly lower concordance for the %pos metric than -MFI and +MFI (Sup. Fig. 12). The comparison between these metric values based on cytoFlagR’s automatically determined threshold and the consensus expert thresholds resulted in similar, or even stronger correlations, across the different controls (Sup. Fig. 13), demonstrating the reliability of cytoFlagR’s automated threshold selection. The user is, however, able to adjust the automatically determined thresholds as desired.

Only a few, specific markers contributed to the minor variability between the metric values based on automated and consensus expert-defined and the highest and lowest expert thresholds (Sup. Fig. 14, 15). More details are provided in the Supplementary results.

### 3.10. Evaluating the impact of sample size on the stability of EMD calculations

By default, cytoFlagR randomly selects 20,000 cells per file for pairwise EMD calculations. The robustness of the EMD measure considering variable numbers of randomly selected cells, from 50 to 50,000 cells, was evaluated. Four example pairs of a sample each from two batches were selected and pairwise EMDs were calculated for each of 10 random sampling iterations. The EMD measures for the spectral dataset (Sup. Fig. 16) and the CytoNorm dataset (Sup. Fig. 17) were stable at 20,000 cells per sample, which established this value as the default parameter in cytoFlagR

## 4 Discussion

High-parameter cytometry datasets are prone to technical aberrations when samples are processed over a period of weeks, months or even years, a phenomenon commonly known as batch effects. While several automated methods exist to correct for batch effects, including CytoNorm (Van Gassen *et al*. 2020b), CytofRUV (Trussart *et al*. 2020), CytofBatchAdjust (Schuyler *et al*. 2019), and cyCombine (Pedersen *et al*. 2022), there are currently no dedicated tools that perform an objective, automated assessment, based on robust statistics, to detect technical problems in batches and markers in cytometry datasets on a per marker basis and within cell populations.

CytoFlagR was developed to address this gap by automatically evaluating cytometry datasets to flag outliers as potentially problematic batches and markers based on several complementary metrics. On a per-marker level, these metrics include assessments of differences in the median intensities of the negative and positive cell populations (-MFI, +MFI) and the percent of positive cells (%pos), using an IQR-based approach. By using these commonly used cytometry measurements, cytoFlagR offers an easily interpretable assessment of potential batch effects in cytometry data. An additional per-marker metric is the evaluation of the overall distributions of fluorescence (for conventional or spectral flow) or signal (for mass cytometry) intensity distributions between batches based on EMDs. EMDs have previously been proposed to identify differences in flow and mass cytometry data between study groups (Orlova *et al*. 2016; Yi and Stanley 2022). Together, these assessments provide users with a comprehensive evaluation of their data, which is presented in detailed and summarizing visualizations.

The detection rate of potential batch effects is influenced by the number of batches present in the dataset. The IQR-based metrics showed a greater decline in the detection rate of problematic batches as the number of batches decreased compared to the EMD-based assessment. This is likely due to the IQR-based metrics relying on measures of the central tendency of the data, which are sensitive to the number of available data points to define outliers. In contrast, the EMD metric is based on pairwise comparisons between batch samples, and the median EMD of each batch is considered its representative value. As a result, the EMD metric is less affected by the number of batches available. This underscores the importance of considering the number of batches when interpreting cytoFlagR’s outputs.

Importantly, marker-level batch effects do not necessarily impact the clustering of control samples into their distinct cell populations. Conversely, some batch effects might only affect specific cell types and may be missed in a global marker-wise assessments (Finak *et al*. 2014; Van Gassen *et al*. 2020b). Therefore, cytoFlagR provides an additional unsupervised clustering-based assessment to evaluate the impact of potentially problematic batches across markers within cell populations. In our analyses, batch-related issues flagged by the IQR-based metrics generally aligned with those determined in the clustered data. However, it is important to consider both marker-wise and cell population-level assessments together when evaluating a dataset for batch effects, and batches that are flagged consistently across multiple controls and metrics indicate more systematic technical issues.

CytoFlagR is primarily designed for assessing datasets based on reference control samples, which are osen included in large scale, long-term cytometry studies. However, recognizing that controls are not always available and that several batch normalization methods do not rely on control samples (Hahne *et al*. 2010ti Pedersen *et al*. 2022ti Quintelier *et al*. 2025), we have also demonstrated cytoFlagR’s ability to evaluate datasets consisting only of biological samples. However, when interpreting the results of cytoFlagR applied to biological samples, it is critical to account for biological variability within the dataset, as the probable batch effects highlighted may reflect biological heterogeneity rather than technical variations in the data.

One limitation of cytoFlagR’s approach is the dependency on the dispersion of metric values and the associated IQR while assessing the positive and negative populations using the IQR-based metrics (-MFI, +MFI, %pos). This can result in either over-flagging, in cases of very low variability between the values for these metrics, or under-flagging, in cases of a large spread of the metric values. The laner case is particularly observed in situations in which a set of batches shows markedly different signal intensity distributions from another set of batches, as seen with CD66 in the CytoNorm dataset (Sup. Fig. 5B). However, this difference in distributions between sets of batches may still be successfully detected by the EMD metric (for example, for CD66 in Sup. Fig. 5C,D). Similarly, cases over-flagged by the IQR-based metrics are osen not flagged by the EMD-based assessment, underscoring the value of cytoFlagR’s complementary assessment methods. Future versions of cytoFlagR could include approaches to determine these specific cases and provide additional information to the users to account for these scenarios. Additionally, cytoFlagR can be extended in the future to include more metrics that assess a list of markers simultaneously in addition to the currently implemented approach based on unsupervised clustering. Such an approach was recently proposed in CytoMDS, which first calculates multi-dimensional EMDs across markers and then creates a low dimensional projection of the results using MDS (Hauchamps *et al*. 2025).

In summary, we propose cytoFlagR as a flexible and objective method to evaluate longitudinal, high-parameter cytometry data to identify potentially problematic batches and markers across control or biological samples. By integrating multiple, complementary forms of assessments, the tool provides researchers with an interpretable and extensive overview of potential problems in their data, which are visualized in various ways. This will help guide users in making informed decisions on excluding problematic batches or markers, or applying an appropriate batch normalization algorithm.

