## Supplementary File for "cytoFlagR: A comprehensive framework to objectively assess high-parameter cytometry data for batch effects"

[1]Department of Pharmacology, Physiology & Neurobiology, University of Cincinnati College of Medicine, Cincinnati, OH, USA
[2]Division of Biomedical Informatics, Cincinnati Children's Hospital Medical Center, Cincinnati, OH, USA
[3]University of Cincinnati College of Medicine, Cincinnati, OH, USA
[4]Division of Immunobiology, Cincinnati Children's Hospital Medical Center, Cincinnati, OH, USA
[5]Department of Biomedical Informatics, University of Cincinnati College of Medicine, Cincinnati, OH, USA
[6]Department of Pediatrics, University of Cincinnati College of Medicine, Cincinnati, OH, USA
[7]Division of Allergy and Immunology, Cincinnati Children's Hospital Medical Center, Cincinnati, OH, USA
[8]Division of Biostatistics and Epidemiology, Cincinnati Children's Hospital Medical Center, Cincinnati, OH, USA
[*] Shruti Eswar and Zachary T. Koenig contributed equally as first authors
[$] Tamara Tilburgs and Sandra Andorf contributed equally as senior authors

**Supplementary information**

### 1 Supplementary methods

#### 1.1 Visual assessments

Several common visualizations are created to provide a global overview of the data for initial data inspection and to visualize dissimilarities among the controls (Fig. 1, box B). The visualizations include: i. Plots of the number of cells per batch per control, ii. Multi-Dimensional Scaling (MDS) plots generated using median expressions of all markers [R package *limma v3.60.6*](Ritchie *et al.* 2015), and iii. Uniform Manifold Approximation and Projection (UMAP) plots [R package *umap v0.2.10.0*] of control samples across batches are generated.

#### 1.2 Automatic calculation of marker thresholds

CytoFlagR automatically determines for each marker of a dataset a threshold to divide the data into positive and negative populations. This threshold is calculated by finding the minimum value of the valley region of the negative and positive peaks in the case of a bimodal distribution (modality determined using R package *LaplacesDemon v16.1.6*). For unimodal distributions, the threshold is set between the 75th and 95th percentile, depending on the proximity to zero. If the 95th quantile is less than 0.5 and the 75th quantile is less than 0.1, the 97th quantile is used as the threshold for that marker, whereas if the 95th quantile is greater than 1.0, the threshold is the 80th quantile value. The 85th quantile is set as the threshold when the 75th quantile is greater than 0.5 and the 95th quantile is less than 1.0. These values are determined across batches and control samples and the resultant median is set as the threshold for that marker.

For every marker, density distributions across batches for each control as well as biaxial dot-plots against a reference marker are visualized with the automatically determined thresholds. The user can adjust the automatically determined threshold for each marker based on these visualizations, and the adjusted thresholds will be used for downstream analyses.

#### 1.3 Earth Mover's Distance (EMD) calculation

Consider two samples for batches *Bx* and *By* of marker *m* and control *ct*. For every marker *m* and control *ct*, EMDs between each pair of batch samples are calculated for binned batch distributions using the R package *emdist [v0.3-3]* with default parameter settings (Eq. 1). An equal number of cells per batch is used to ensure reliability of the distance measure (default: 20,000 cells).

$$EMD_{[m,ct,Bx,By]} = EMD[distr(Bx), distr(By)] \quad \text{(Eq. 1)}$$

The EMD determined between two batches can be understood as the amount of effort required to change the shape of the distribution of *Bx* to be equivalent to the shape of *By*. If the distribution *Bx* is considerably different from *By*, their resultant pairwise EMD will be high.

In order to calculate the fourth assessment metric per batch, for each marker and control, the median of the pairwise EMD values is determined for each batch (Eq. 2). This is calculated for all *n* batches present in the dataset. A batch with a median pairwise EMD greater than or equal to 5.0 (default) is considered as a probable batch effect and is flagged.

$$medEMD_{[m,ct,Bx]} = median[EMD_{Bx,By}, \dots ,EMD_{Bx,Bn}] \quad \text{(Eq. 2)}$$

#### 1.4 Evaluations

##### 1.4.1 Analyzing the influence of number of batches on cytoFlagR metrics

To effectively detect potential batch effects in a dataset, it is imperative to evaluate how the number of batches impacts each of cytoFlagR's metrics. This was evaluated for each of the four metrics of cytoFlagR through several steps: First a list of all flags, i.e., all problematic batches that are flagged for a marker and control (defined as a 'flagged case'), in a dataset was generated. Then, for each 'flagged case', each batch was randomly eliminated one at a time (while retaining the problematic batch) until only two batches remained. This was repeated 50 times for every 'flagged case' in every metric. For every specific number of batches (*Nb*) considered, the number of iterations that a 'flagged case' was successfully detected as a 'flag' was determined and divided by 50 (the total number of iterations), to calculate the detection rate (Eq. 3).

$$Detection\ Rate\ of\ flagged\ case_{[Nb]} = \frac{Number\ of\ iterations\ with\ flagged\ case\ detected\ [for\ Nb\ batches]}{Total\ number\ of\ iterations} * 100$$

(Eq. 3)

Per metric, a confidence interval (CI) of the detection rate for all 'flagged cases' was estimated and visualized along the mean detection rate as a summary line plot.

##### 1.4.2 Assessing the impact of variable thresholds for separating negative and positive populations on the IQR-based metrics

To assess the variability among researchers in setting thresholds to separate cells into positive and negative populations for a marker, five researchers experienced in cytometry analysis independently determined thresholds for each marker in the spectral dataset. The lowest and highest expert defined thresholds were compared for each of the IQR-based metrics (-MFI, +MFI, %pos), marker and control sample.

Additionally, the thresholds determined automatically by cytoFlagR were compared to the consensus expert-defined threshold values. These comparisons were visualized using dot-plots of -MFI, +MFI and %pos values of all markers along with the identity lines and Pearson correlation coefficients for each control.

##### 1.4.3 Quantifying the effect of sample size on robustness of EMD calculations

Since the EMD assessment requires calculating EMDs for each pair of batch samples for each control and marker in a dataset, cytoFlagR down-samples the number of cells per sample. By default, it randomly selects 20,000 cells per batch per control to establish pairwise EMDs for a marker and control. To assess the impact of the number of randomly selected cells per sample on the robustness of the EMD measure, EMDs between a pair of batches are calculated across different numbers of randomly selected cells, ranging from 50 to 50,000 cells. Each random sampling was performed ten times and the calculated EMDs are visualized with boxplots for each of the different sample sizes.

#### 1.5 Dataset description

One of the datasets used for testing cytoFlagR is the CytoNorm CyTOF dataset (Van Gassen *et al.* 2020a), consisting of 10 batches of two unstimulated and two Interferon-alpha and Lipopolysaccharide (IFNa-LPS) stimulated control samples of whole blood from healthy donors with 32 protein channels. This dataset is frequently utilized to test different batch normalization methods (Trussart *et al.* 2020; Van Gassen *et al.* 2020a; Pedersen *et al.* 2022).

cytoFlagR was also applied to four control samples from an original spectral cytometry dataset. Prepared samples for 22 protein channels were analyzed across 18 different days, each considered a batch. Details are described below.

Since some studies do not include reference control samples, mass cytometry FCS files of biological samples from the HealthNuts (Neeland *et al.* 2020; Tursi *et al.* 2022) study were used to test the ability of cytoFlagR to highlighted probable batch effects in a dataset without reference controls. Unstimulated (US), phorbol 12-myristate 13-acetate ionomycin stimulated (PMA) and peanut-protein (PN) stimulated PBMC samples with 22 protein channels were analyzed across 6 different days (batches). Samples from each of the three study groups (infants with peanut sensitization but tolerance to peanut, infants that were clinically peanut allergic, and non-sensitized, non-peanut allergic infants) were included in each of the batches. The gating strategy for selection for live single cells was presented in our original publication (Neeland *et al.* 2020).

#### 1.6 Spectral flow cytometry dataset: experimental details

Technical controls were obtained from healthy, non-pregnant adult peripheral blood donors at a local blood donation center. Peripheral leukocytes were isolated by Ficoll gradient centrifugation for 20 min at 800g. Leukocytes from each donor were washed once with wash medium (RPMI, 10% FBS, pen/strep) and centrifugated for 7min at 650g. Unstimulated controls 1 and 2 were directly aliquoted in X-VIVO supplemented with 10% FBS and 10% DMSO and frozen in liquid nitrogen. Stimulated controls 3 and 4 were cultured in X-VIVO medium supplemented with 5% human serum and 50 U/mL IL-2 in the presence of anti-CD3/CD28 beads (BD Biosciences 1ug/ml) for 7 days. Leukocytes were collected on day 7 post-stimulation, aliquotted and frozen in liquid nitrogen as described. Controls were thawed at 37°C, washed once and directly stained for analysis on a Cytek Aurora spectral flow cytometer. For surface staining, cells were stained for 30 min in RPMI containing 1% FBS and pen/strep. For intracellular staining, cells were fixed and permeabilized using the eBiosciences FOXP3 kit (eBiosciences). All flow cytometric data were acquired using equipment maintained by the Research Flow Cytometry Core in the Division of Rheumatology at Cincinnati Children's Hospital Medical Center. Initial data processing and gating on CD45+CD14- lymphocytes was performed using FlowJo software [*v10.7.2*] and data were exported as FCS files.

##### 1.7 Data availability and software implementation

The CytoNorm dataset is publicly available on FlowRepository under ID FR-FCM-Z247 (Spidlen *et al.* 2012; Van Gassen *et al.* 2020b). The HealthNuts dataset is available on ImmPort under ID SDY215 (Bhattacharya *et al.* 2014, 2018; Tursi *et al.* 2022). The raw data of the spectral cytometry dataset can be obtained from Zenodo under https://zenodo.org/uploads/15388817 (doi: 10.5281/zenodo.15388817) (European Organization For Nuclear Research; OpenAIRE 2013)

##### 1.8. Software implementation

All scripts were compiled using the R programming language (version 4.4.1). cytoFlagR is open-source and available with documentation and an example on GitHub (https://github.com/AndorfLab/cytoFlagR).

#### 2 Supplementary results

##### 2.1 Results of the application of CytoFlagR to the CytoNorm dataset

Applying cytoFlagR to CD25 in the CytoNorm dataset showed that PTLG025 and PTLG021 were consistently flagged as potentially problematic batches across the -MFI, +MFI and %pos metrics (Sup. Fig. 4B) for control_2 as well as in the pairwise and median EMDs for control_1, control_2, and control_4 (median EMD values ≥ 3, default threshold for mass cytometry dataset controls) (Sup. Fig. 4C). Control_2 showed the overall greatest median EMDs for PTLG025 and PTLG021 (Sup. Fig. 4D).

For CD66, one set of batches (PTLG021, PTLG025, PTLG026, and PTLG027) had visually distinct marker expression density distributions when compared with the set of the 6 remaining batches. As a result, the IQR-based metrics showed a wide spread of values, with one set of the batches showing values for each of the IQR-based metrics on one end of the range and the other set of batches on the other end (Sup. Fig. 5B). Consequently, none of the batches were flagged in these metrics. However, one set of batches (PTLG021, PTLG025, PTLG026, PTLG027) was highlighted as having potential technical issues across all four controls by the EMD metric, since these batches had greater degrees of dissimilarities to all batches of the other set of batches (Sup. Fig. 5 C, D).

The outcome of cytoFlagR assessments on all markers present in the CytoNorm dataset was summarized across controls, batches and markers and visualized as a heatmap (Sup. Fig. 6). Batches PTLG033, PTLG026, PTLG034 and PTLG021 ranked at the top with the highest number of markers flagged across controls along with the Inhibitor of Nuclear Factor Kappa B (IkB) marker which had the greatest number of flagged batches across controls.

The results of unsupervised clustering of the control samples of CytoNorm were examined next. The control samples were clustered into 25 clusters using FlowSOM based on 21 phenotypic markers (Sup. Fig. 7A). Batch PTLG021 in Cluster 3, PTLG034 in Cluster 17, PTLG034 in Cluster 21, and PTLG031 in Cluster 22 showed higher proportions of cells of a batch than expected, highlighting them as possibly problematic batches. Specifically, Cluster 22 had a greater than expected proportion of batch PTLG031 in control 4 (Sup. Fig. 7B, top). Clusters with an even distribution of batches across controls were also depicted (Sup. Fig. 7B, bottom). A UMAP projection of the clustered data (Sup. Fig. 7C) supported the differences in batch contributions within the different clusters.

##### 2.2 The impact of inter-expert variability of marker thresholds on IQR-based metrics and comparison with automated marker threshold determination

To assess the impact of this variability on cytoFlagR's IQR-based metrics, -MFI, +MFI and %pos metric were calculated using the lowest and highest defined threshold values among five experts for each marker and control sample in the spectral dataset (Sup. Fig. 12). The MFI metric values showed strong correlation between those derived based on the lowest and highest expert thresholds, with all Pearson correlation coefficients > 0.944 for -MFI and > 0.962 for +MFI across the 4 controls (Sup. Fig. 12 A, B). Inter-expert variability was more evident in the %pos metric (Sup. Fig. 12C) with Pearson correlation coefficients ranging from 0.839 to 0.981 across controls, indicative of a degree of variability in setting these thresholds among researchers.

The performance of the thresholds that were automatically determined by cytoFlagR was evaluated using a similar approach, comparing values for the IQR-based metric values derived using the automated thresholds with those based on the consensus expert-defined thresholds (Sup. Fig. 13). All three IQR-based metrics showed a large correlation between values based on the automated and consensus expert thresholds (all Pearson correlation coefficients > 0.91).

The overall equal or greater correlation between IQR-metric values based on the automatically determined and consensus expert thresholds compared to between the lowest and highest expert thresholds demonstrate the reliability of cytoFlagR's automated threshold selection.

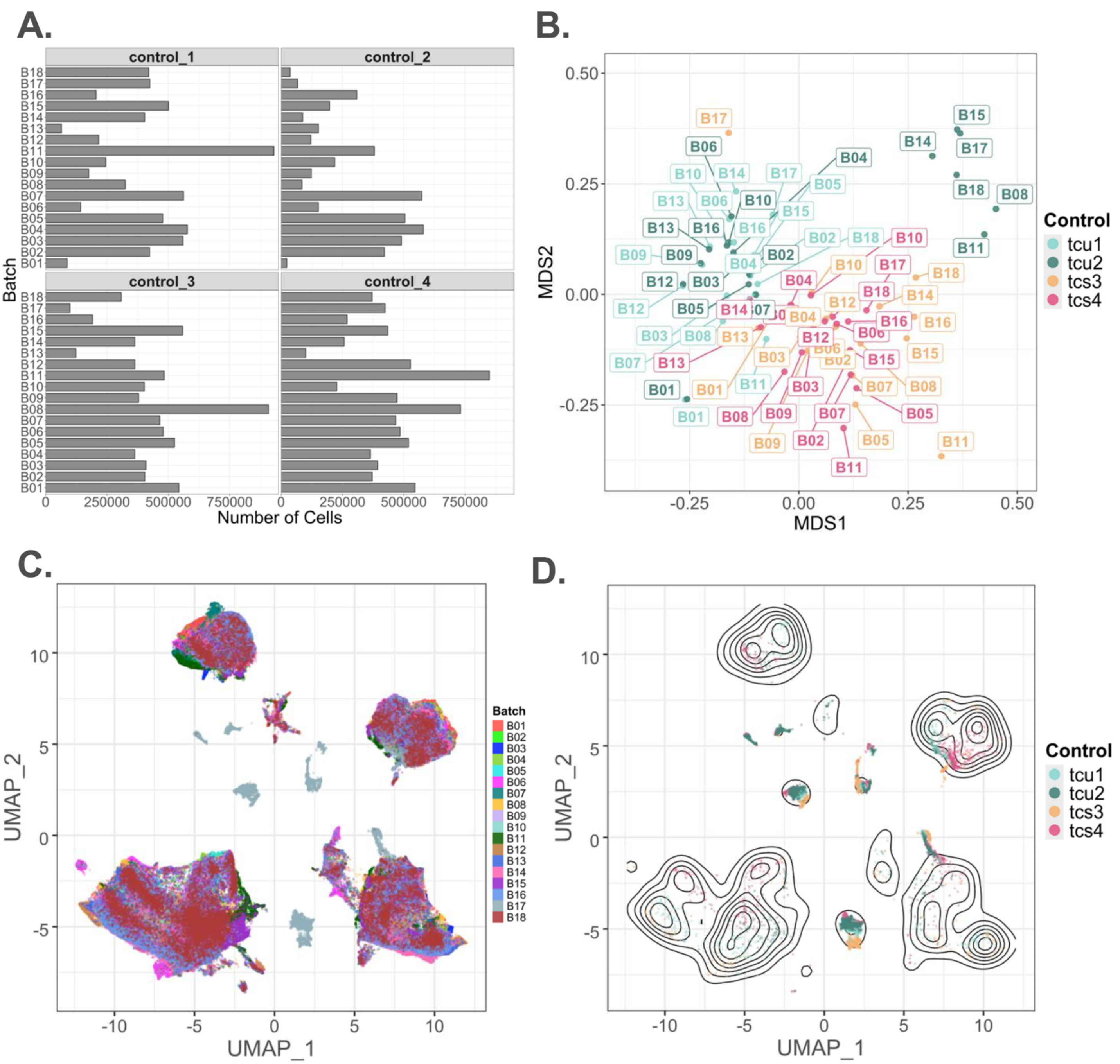


**Supplementary Figure 1. Initial, exploratory examination of spectral dataset. (A)** Bar plot of number of cells present in each control sample across 18 batches. **(B)** Multi-dimensional Scaling (MDS) plot of data annotated by batch labels and colored by control samples to visualize inter-control discrepancies. **(C)** A Uniform Manifold Approximation and Projection (UMAP) of the data subsampled to 4,000 cells per samples colored by batch to check for even distribution of cells by batch. The data shows batch B17 separated from the other batches. **(D)** Cells from batch B17 are visualized as dots and cells from all other batches by contour plots, further highlighting a potential technical effect impacting this batch.

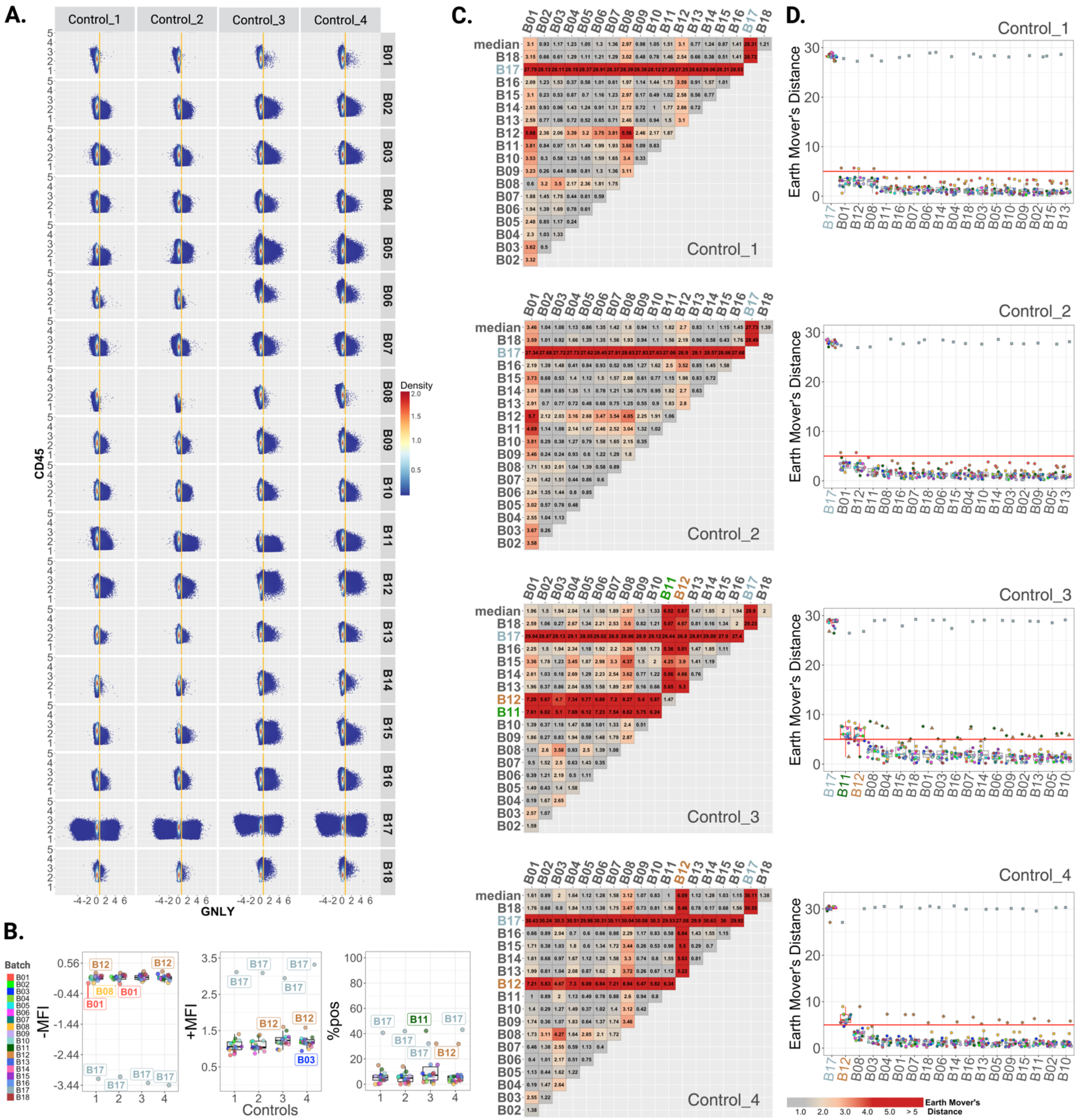


**Supplementary Figure 2. cytoFlagR detects potentially problematic batches across 4 controls in GNLY in the spectral dataset. (A)** Biaxial dot-plots of GNLY vs. CD45 across 18 batches of two unstimulated (control_1, control_2) and two stimulated (control_3, control_4) control samples. Yellow lines indicate the threshold at 0.7 separating the negative and positive populations. **(B)** Batch B17 is a consistent flag in all three metrics, with B01 flagged in the unstimulated controls for -MFI, along with B12 in controls 1 and 4, as well as B08 in control_1. The stimulated controls also highlight that B12 and B03 are flagged in control_4 in the +MFI metric. B01 and B08 contained <100 cells in the positive populations and were excluded from the +MFI metric assessment. The %pos metric detects B11 in control_3 and B12 in control_4 as potentially problematic batches. **(C)** Pairwise and median EMDs across batches for each control visualized with heatmaps. **(D)** EMDs ordered by their respective median values for each batch are visualized as boxplots for each control, highlighting B17 in all four control samples as a potentially problematic batch, B12 in the stimulated controls, along with B11 in control 3 (represented by boxplots outlined in red). These batches show a median EMD greater than the default threshold of 5 (red horizontal lines).

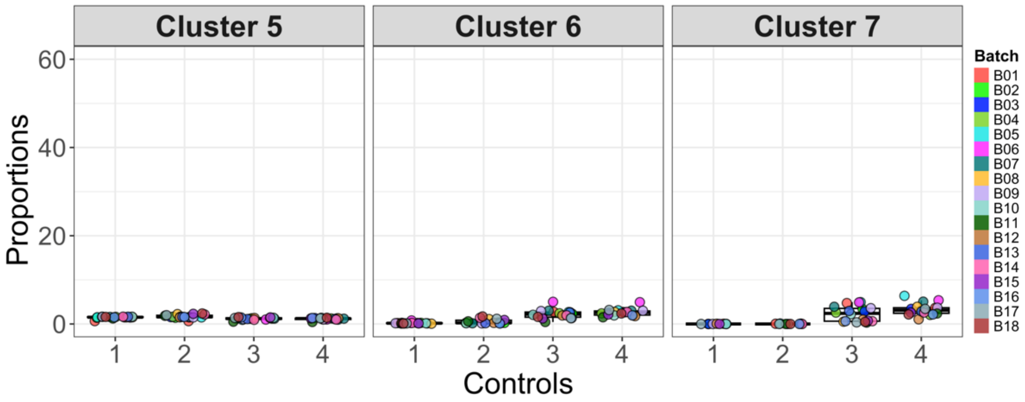


**Supplementary Figure 3.** Batch proportions present in each cluster (not visualized in Figure 5B), across all four control samples are represented by boxplots. The visualized clusters have an even distribution of batches, suggesting that they are not impacted by technical variations in the control samples.

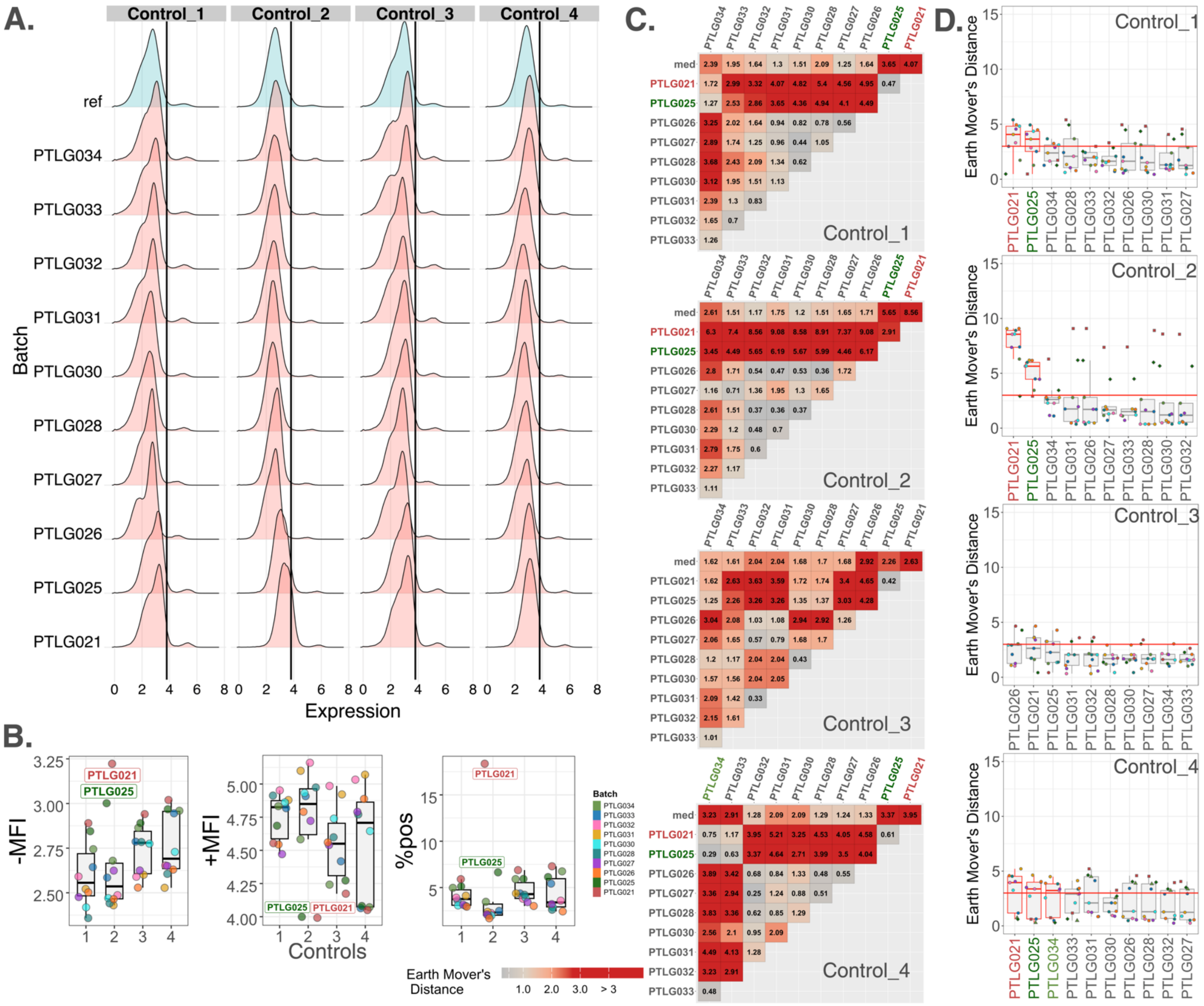

**Supplementary Figure 4. cytoFlagR detects potentially problematic batches across 4 controls in CD25 of the CytoNorm dataset. (A)** Density distributions for CD25 across 10 batches of two unstimulated (control_1, control_2) and two stimulated (control_3, control_4) control samples. The black line at 3.8 indicates the threshold separating negative and positive populations. **(B)** Batches PTLG025 and PTLG021 have noticeable positive shifts and are consistently highlighted as potentially problematic batches in control_2 by the -MFI metric (left), +MFI metric (middle) and the %pos metric (right). **(C)** Pairwise and median EMDs (med) across batches for each control visualized with heatmaps. **(D)** Median EMDs for each batch ordered by their respective median values are visualized as boxplots for each control. Batches PTLG025 and PTLG021 are detected as potential batch effects in controls 1, 2 and 4 along with batch PTLG034 in control_4. These batches show a median EMD greater than or equal to the threshold of 3 set for mass cytometry datasets (red horizontal lines).

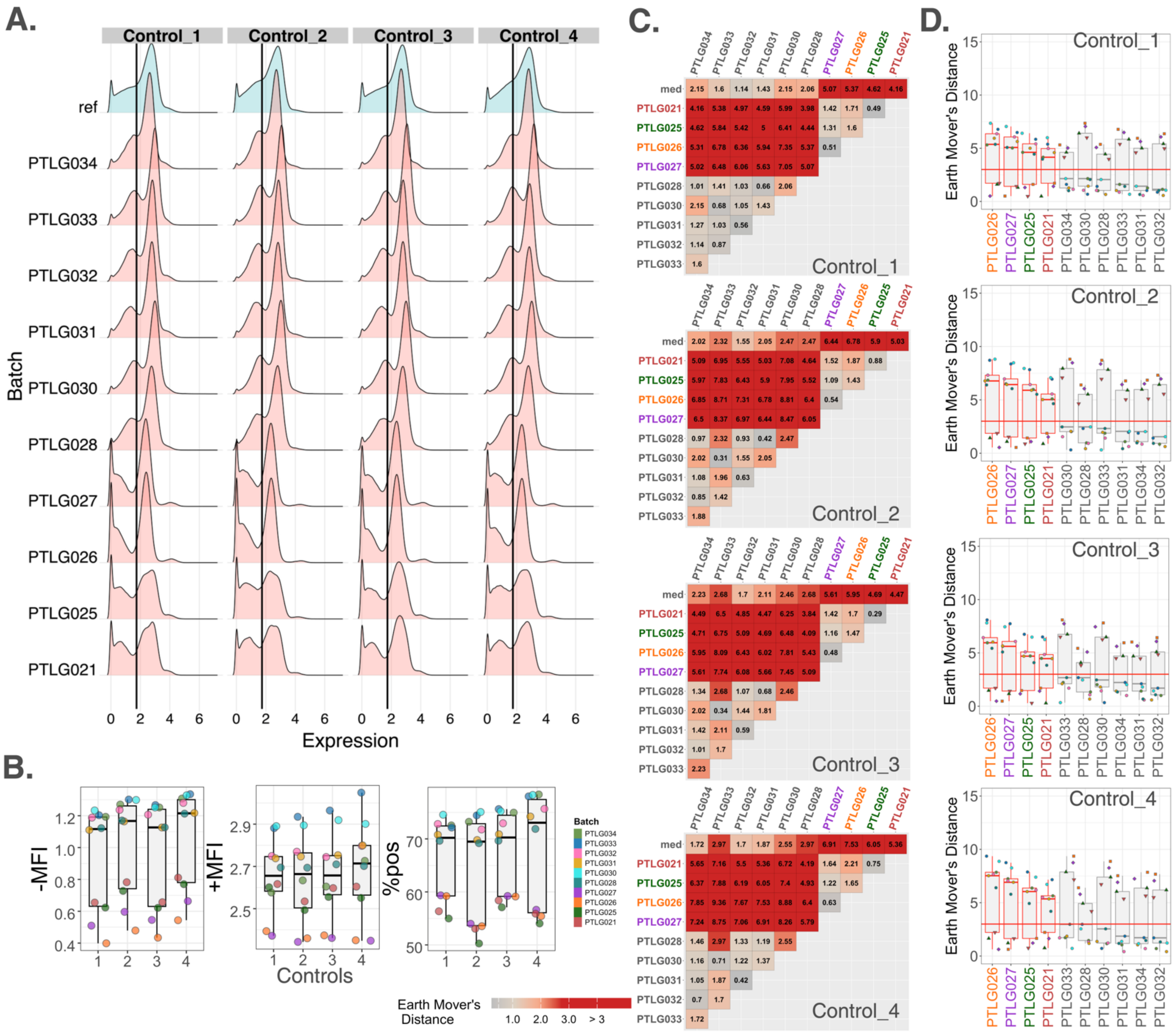


**Supplementary Figure 5. cytoFlagR detects potentially problematic batches across 4 controls in CD66 of the CytoNorm dataset. (A)** Density distributions for CD66 across 10 batches of two unstimulated (control_1, control_2) and two stimulated (control_3, control_4) control samples. The black line indicates the threshold (1.74) separating negative and positive populations. **(B)** Boxplots of values of the -MFI (left), +MFI (middle) and %pos (right) for all batch samples stratified by controls. **(C)** Pairwise and median EMDs across batch samples for each control visualized with heatmaps and **(D)** the median EMDs shown as boxplots per batch ordered by their median. The density distributions of batches PTLG027 to PTLG021 (bottom 4 in **A**) are evidently different from those of batches PTLG034 through PTLG028 (top 6 in **A**), and as a result every control indicates batches PTLG027, PTLG026, PTLG025, and PTLG021 having median EMDs ≥ 3, the threshold set for mass cytometry datasets (red horizontal lines), flagging them as potentially showing a batch effect.

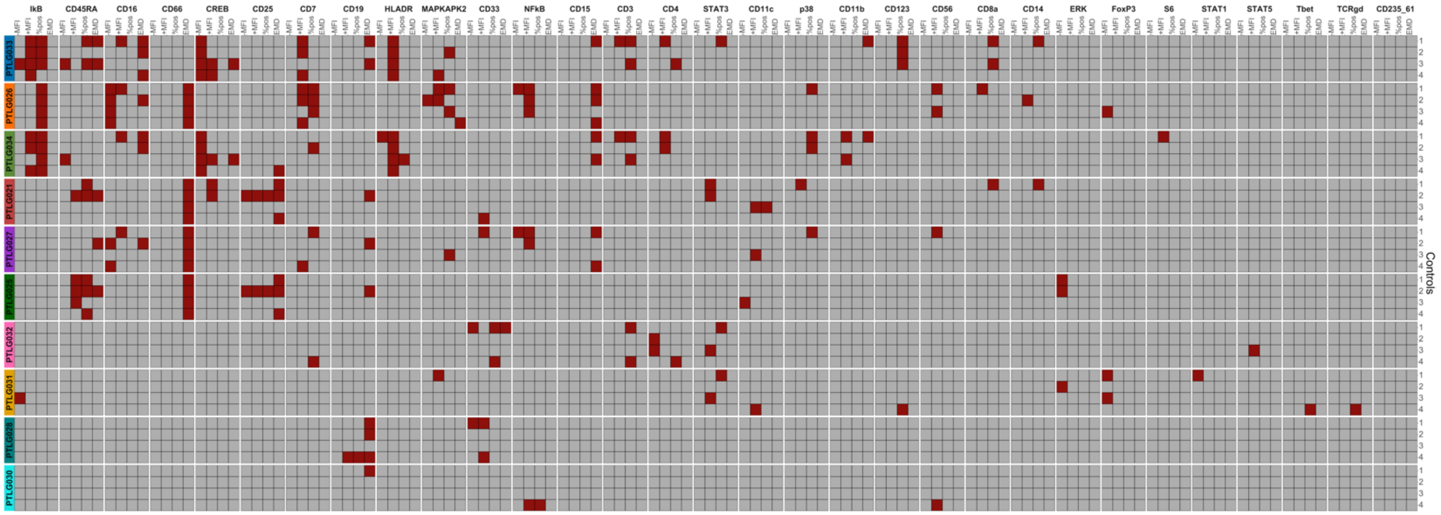


**Supplementary Figure 6. Heatmap summarizing potential batch effects flagged across markers, controls, and batches in the CytoNorm mass cytometry dataset.** The CytoNorm dataset consisting of 4 controls, 10 batches and 31 protein channels was assessed using cytoFlagR to identify potential batch effects across four metrics [-MFI, +MFI, %pos, and EMD (≥ 3)]. Potential batch effects identified for each marker, control, and batch are summarized and visualized with a heatmap, ranking batches and markers from most frequently flagged to least. Dark red tiles indicate flagged potential batch effects and gray tiles indicate no detected batch effects. Batches PTLG033, PTLG026, and PTLG034 show the most probable batch effects across markers and controls.

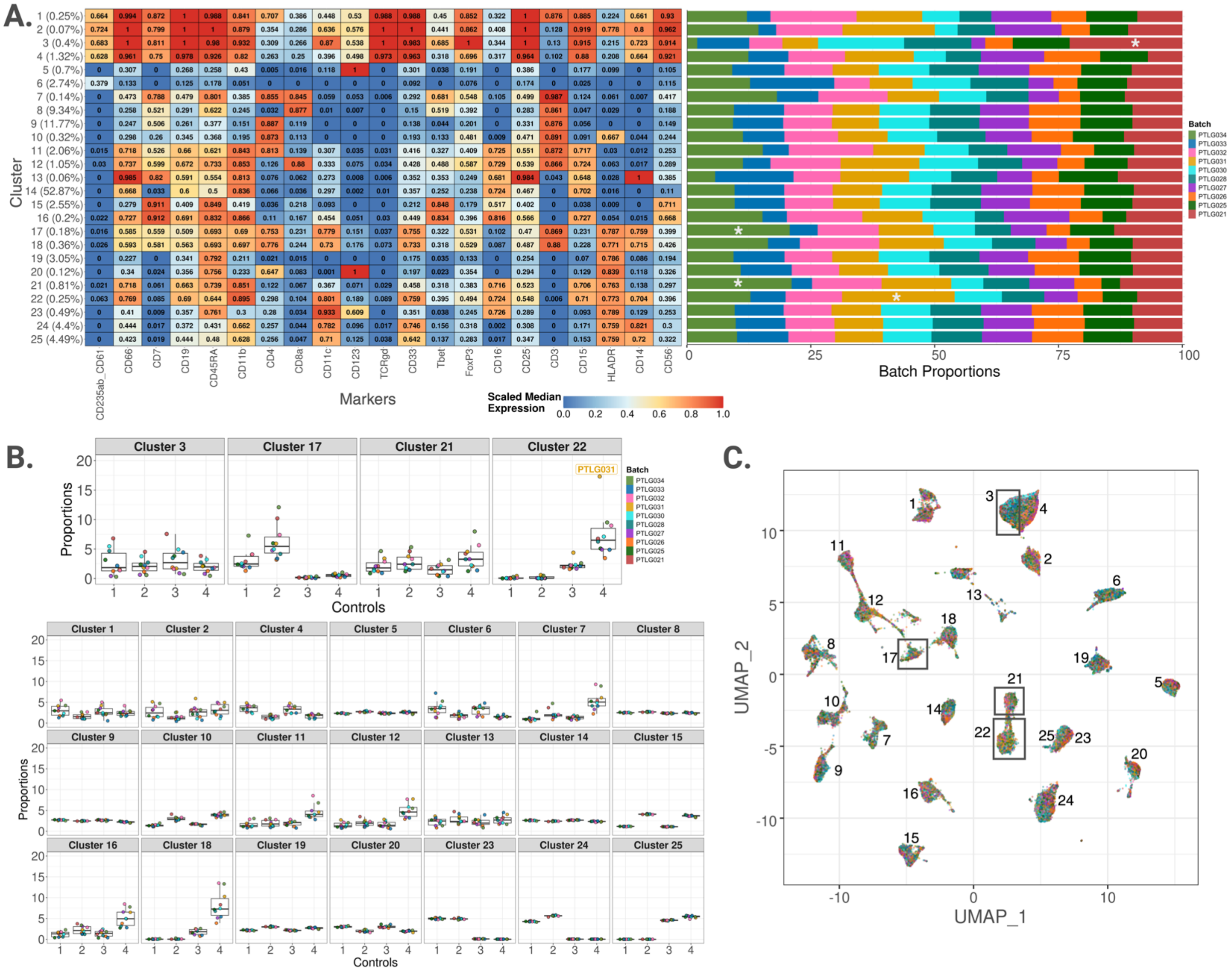


**Supplementary Figure 7. Flagging problematic batches based on unsupervised clustering of the CytoNorm dataset. (A)** Heatmap of marker expressions across 21 markers and 25 clusters (left) and the batch proportions of analyzed cells in each cluster represented by a bar plot (right). Cluster 3 shows a higher-than-expected abundance of PTLG021, Cluster 17 has a greater proportion of PTLG034, Cluster 21 a higher abundance of PTLG034 and Cluster 22 has a higher-than-expected proportion of PTLG031 (marked by *). **(B)** Boxplots representing the batch proportions of these clusters (top row) stratified by controls. The bottom rows illustrated the distribution of cells across controls in the remaining clusters devoid of problematic batches. **(C)** A UMAP of all clusters depicted with analyzed cells colored by batch. The flagged batches highlighted by black boxes corroborates the results of (A) and (B).

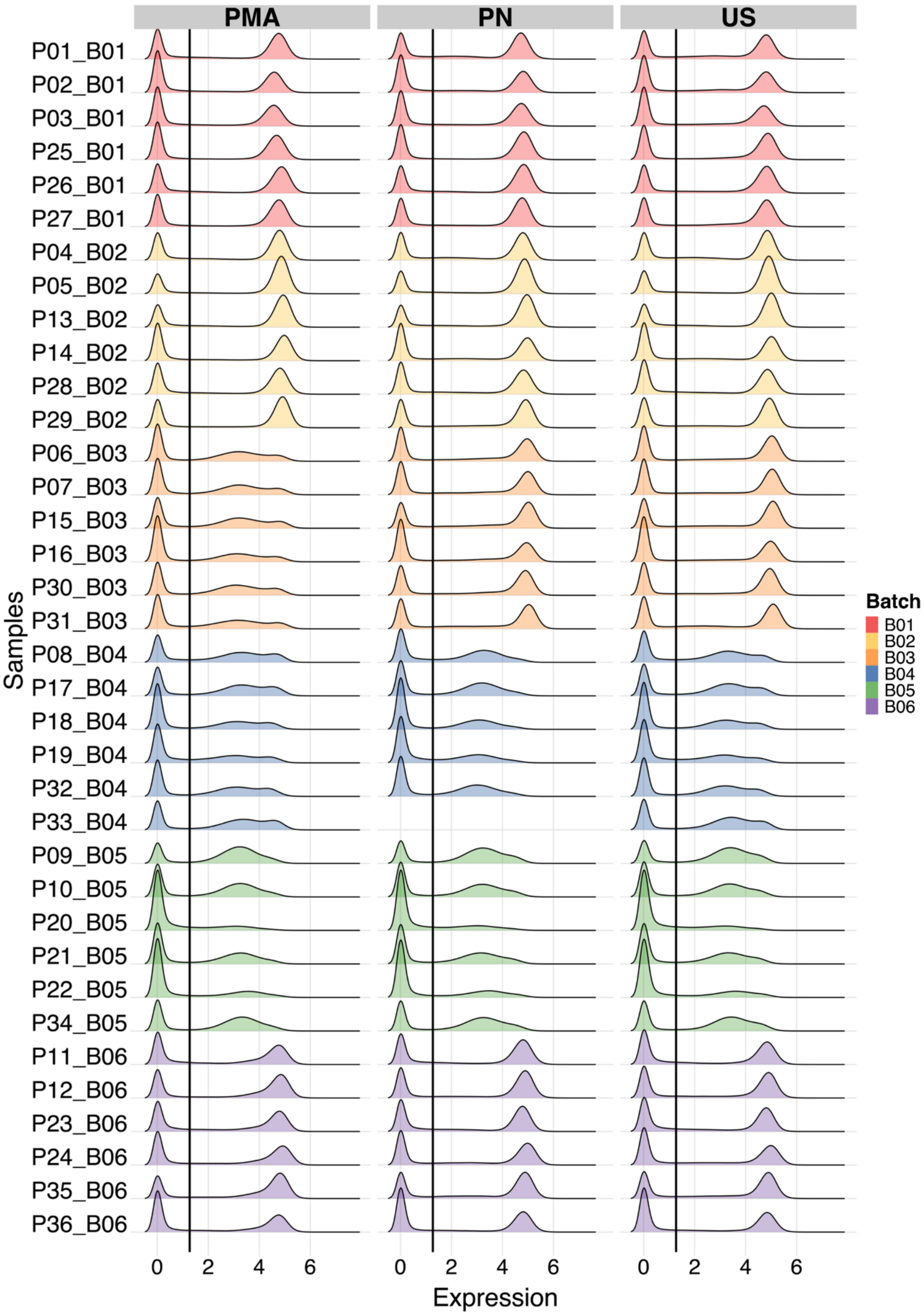


**Supplementary Figure 8. Density distributions of CD27 across 36 samples within 6 batches for 3 stimulations** – phorbol 12-myristate 13-acetate ionomycin (PMA) stimulation, peanut protein solution (PN) stimulation, and unstimulated (US) – from the HealthNuts dataset. The solid black line indicates the threshold to divide the data into positive and negative populations. Sample P33 was not available under PN stimulation. Densities of all other markers for US samples were previously published (Tursi *et al.* 2022).

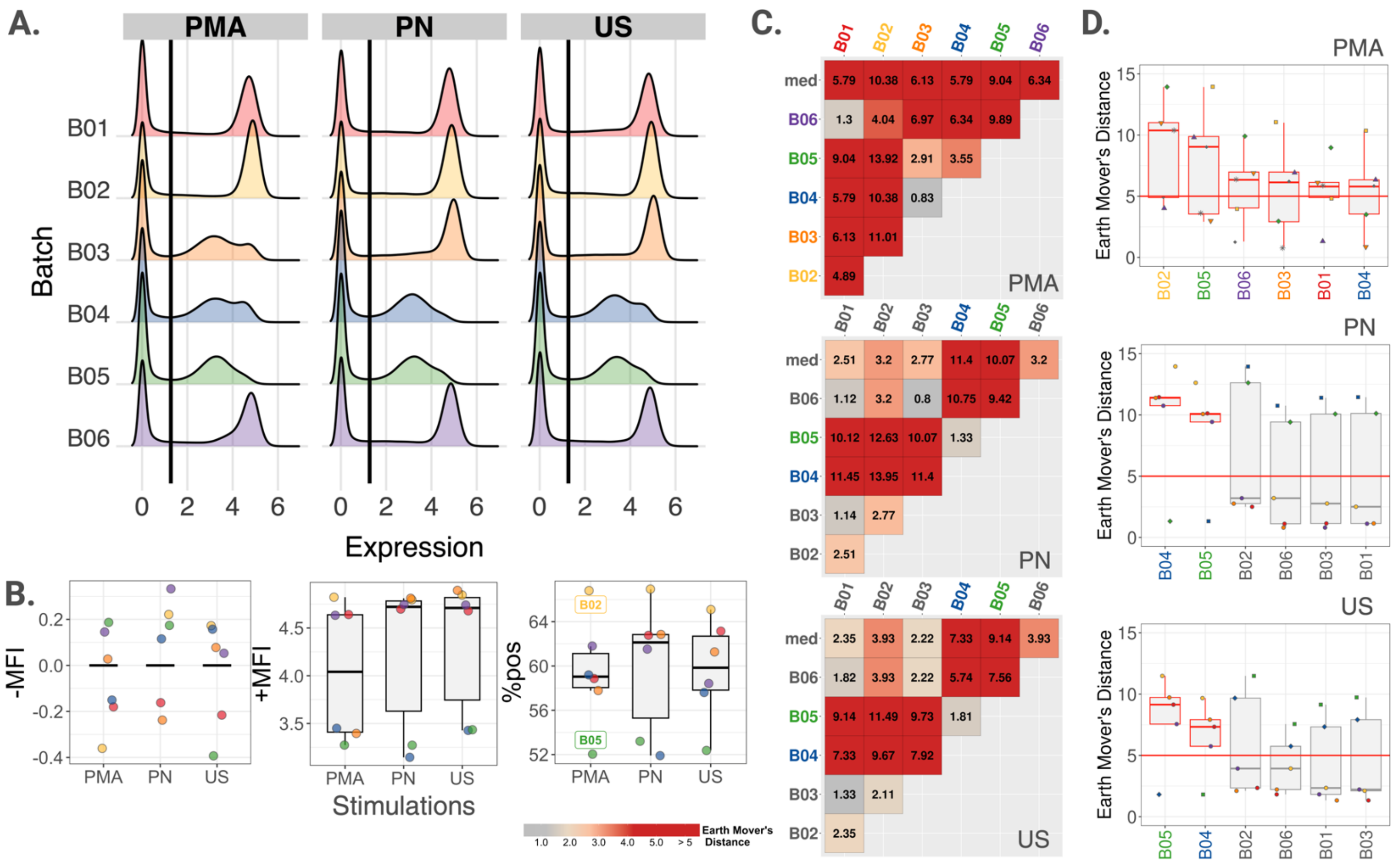

**Supplementary Figure 9. cytoFlagR detects potentially problematic batches across 3 stimulation statuses in CD27 of the HealthNuts dataset.** 45,000 cells per sample were randomly selected and concatenated per batch to generate a sample-balanced pooled distribution per batch. **(A)** Pooled density distributions for CD27 across 6 batches for phorbol 12-myristate 13-acetate ionomycin stimulated (PMA), peanut protein solution stimulated (PN), and unstimulated (US) samples. The black line at 1.27 indicates the threshold separating negative and positive populations. **(B)** Boxplots of distribution of batches for the -MFI (left), +MFI (middle), and %pos metric (right) of CD27 stratified by stimulation. B02 and B05 were flagged as exhibiting probable batch effects in the %pos metric for the PMA stimulation. **(C)** Pairwise and median EMDs across batches for each stimulation visualized with heatmaps. **(D)** EMDs ordered by their respective median values for each batch are visualized as boxplots for each stimulation indicate all batches as highly dissimilar overall for the PMA stimulation, along with B04 and B05 for the PN and US stimulation with their respective median EMD values ≥ 5 (red horizontal lines).

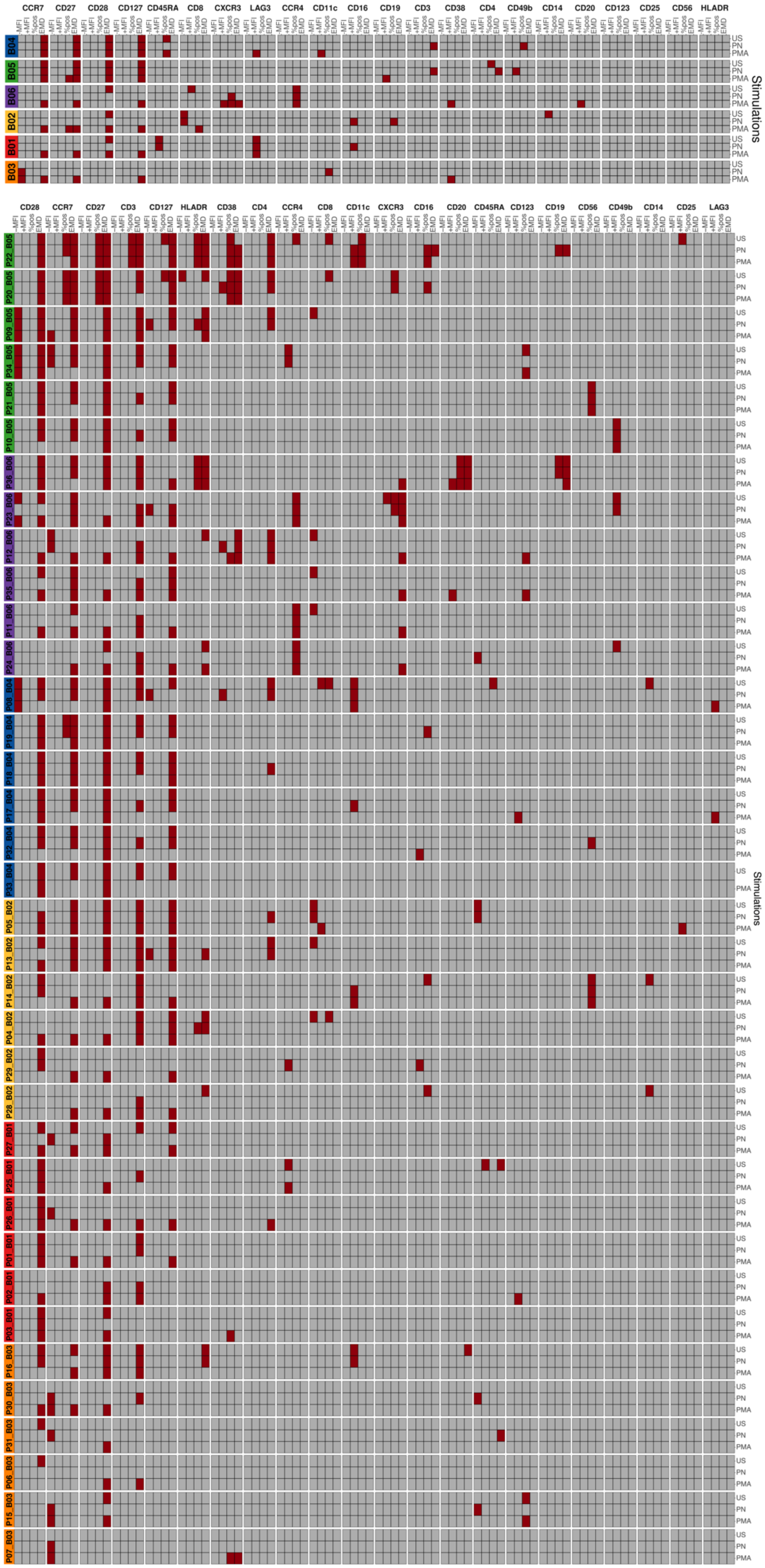
CCR7 CD27 CD28 CD127 CD45RA CD8 CXCR3 LAG3 CCR4 CD11c CD16 CD19 CD3 CD38 CD4 CD49b CD14 CD20 CD123 CD25 CD56 HLADR
-MFI +MFI %pos EMD
B04
B05
B06
B02
B01
B03
US PN PMA
Stimulations
CD28 CCR7 CD27 CD3 CD127 HLADR CD38 CD4 CCR4 CD8 CD11c CXCR3 CD16 CD20 CD45RA CD123 CD19 CD56 CD49b CD14 CD25 LAG3
-MFI +MFI %pos EMD
P22 B05
P20 B05
P09 B05
P34 B05
P21 B05
P10 B05
P36 B06
P23 B06
P12 B06
P35 B06
P11 B06
P24 B06
P08 B04
P19 B04
P18 B04
P17 B04
P32 B04
P33 B04
P05 B02
P13 B02
P14 B02
P04 B02
P29 B02
P28 B02
P27 B01
P25 B01
P26 B01
P01 B01
P02 B01
P03 B01
P16 B03
P30 B03
P31 B03
P06 B03
P15 B03
P07 B03
US PN PMA
Stimulations

**Supplementary Figure 10. Heatmap summarizing potential batch effects flagged across markers, stimulations, samples, and batches in the HealthNuts dataset.** The HealthNuts dataset was assessed utilizing pooled samples per batch (top) and on a per file basis (bottom) using cytoFlagR to identify potential batch effects across four metrics (-MFI, +MFI, %pos, and EMD). Potential batch effects identified for each marker, stimulation, and batch across samples are summarized and visualized with a heatmap, ranking batches and markers from most frequently flagged to least. Dark red tiles indicate flagged potential batch effects and gray tiles indicate no detected batch effects. Samples P22 and P20 of batch B05 as well as P36 and P23 of batch B06 show the most probable batch effects across markers and stimulations.

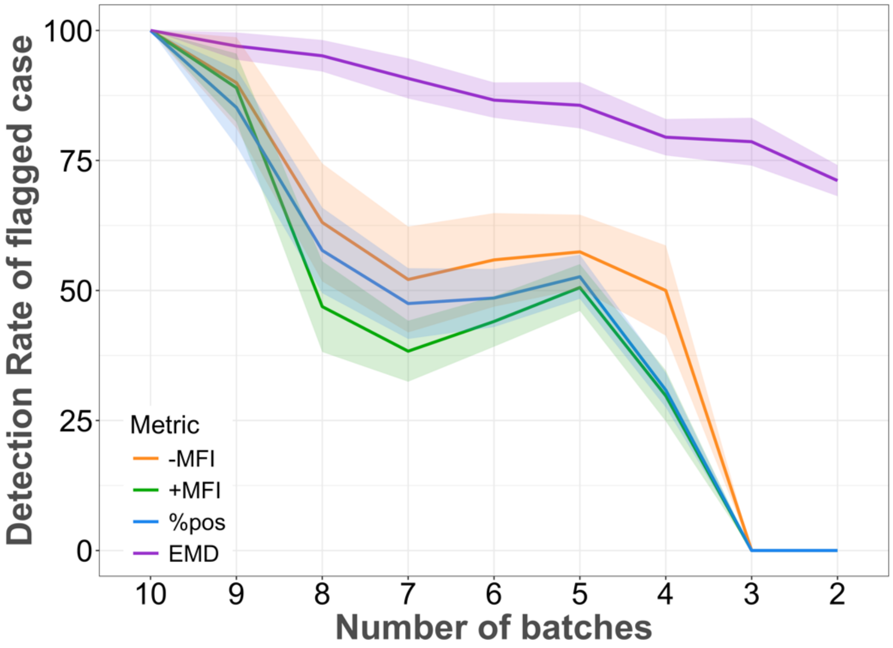


**Supplementary Figure 11. Impact of number of batches used for the four cytoFlagR assessment metrics (CytoNorm dataset).** The lines represent the mean detection rate of flagged cases using different number of batches for each of the four metrics [-MFI, n = 32 (total flagged cases of the entire dataset of 10 batches); +MFI, n =49; %pos, n = 49; EMD, n = 46]. 95% confidence intervals based on 50 repeats of successively reducing number of batches at random are shown as shaded areas. The cases flagged by -MFI, +MFI, and %pos metrics exhibit a downward trend, concluding in an abrupt decline to zero. In contrast, the EMD metric shows a consistently decreasing trend thereby highlighting the influence of number of batches has on the assessment metric.

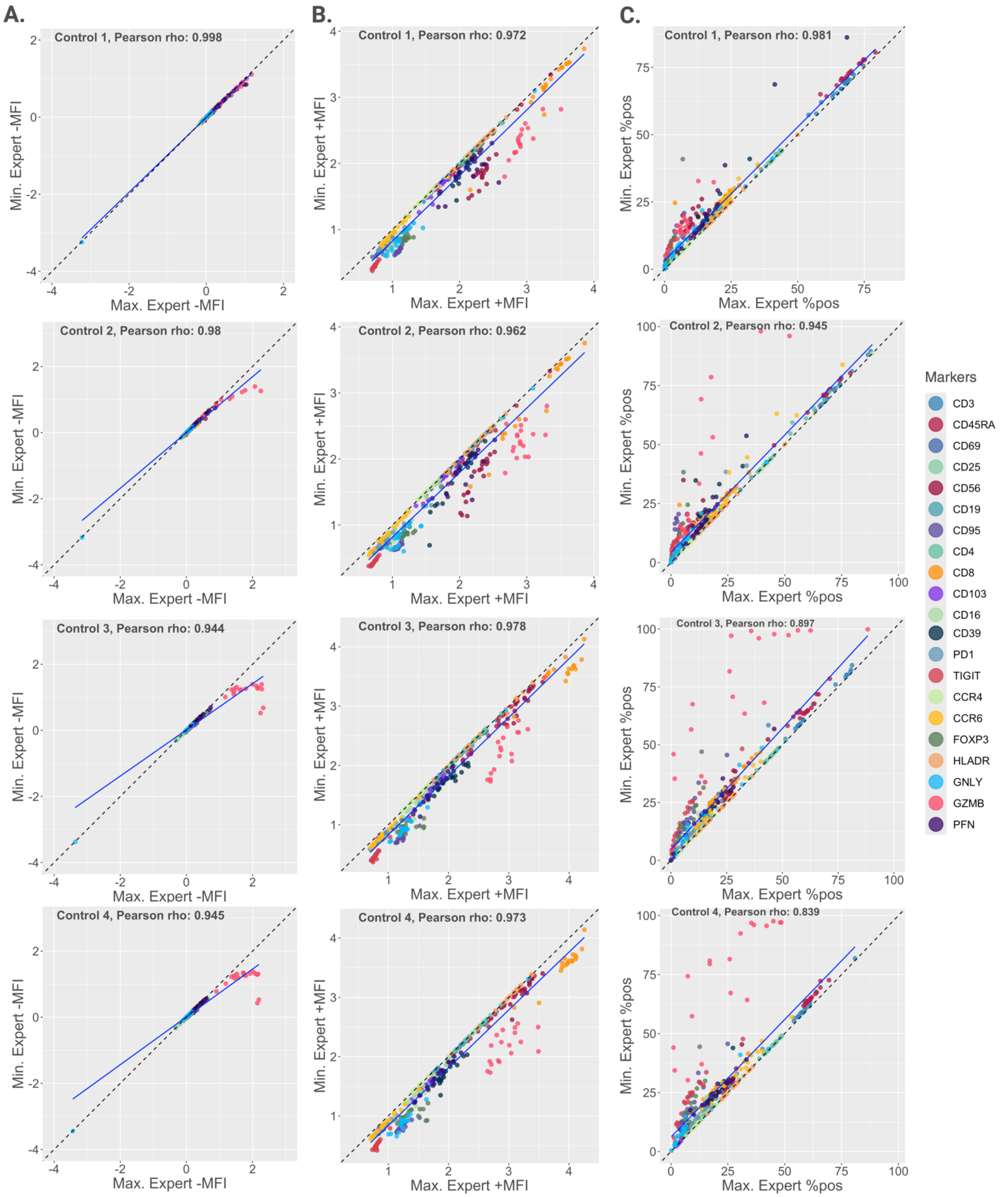


**Supplementary Figure 12. Variability of IQR-based metrics based on thresholds for separating negative and positive populations set by five experts (spectral dataset).** The lowest and highest threshold among 5 experts were used to calculate the three metrics **(A)** -MFI, **(B)** +MFI and **(C)** %pos for each marker and each control sample. The resulting values are represented using dot-plots (x-axis result based on the largest threshold, y-axis result based on the lowest threshold). Pearson's correlation coefficients (r) and their corresponding correlation lines are illustrated. The dashed line represents the identity line.

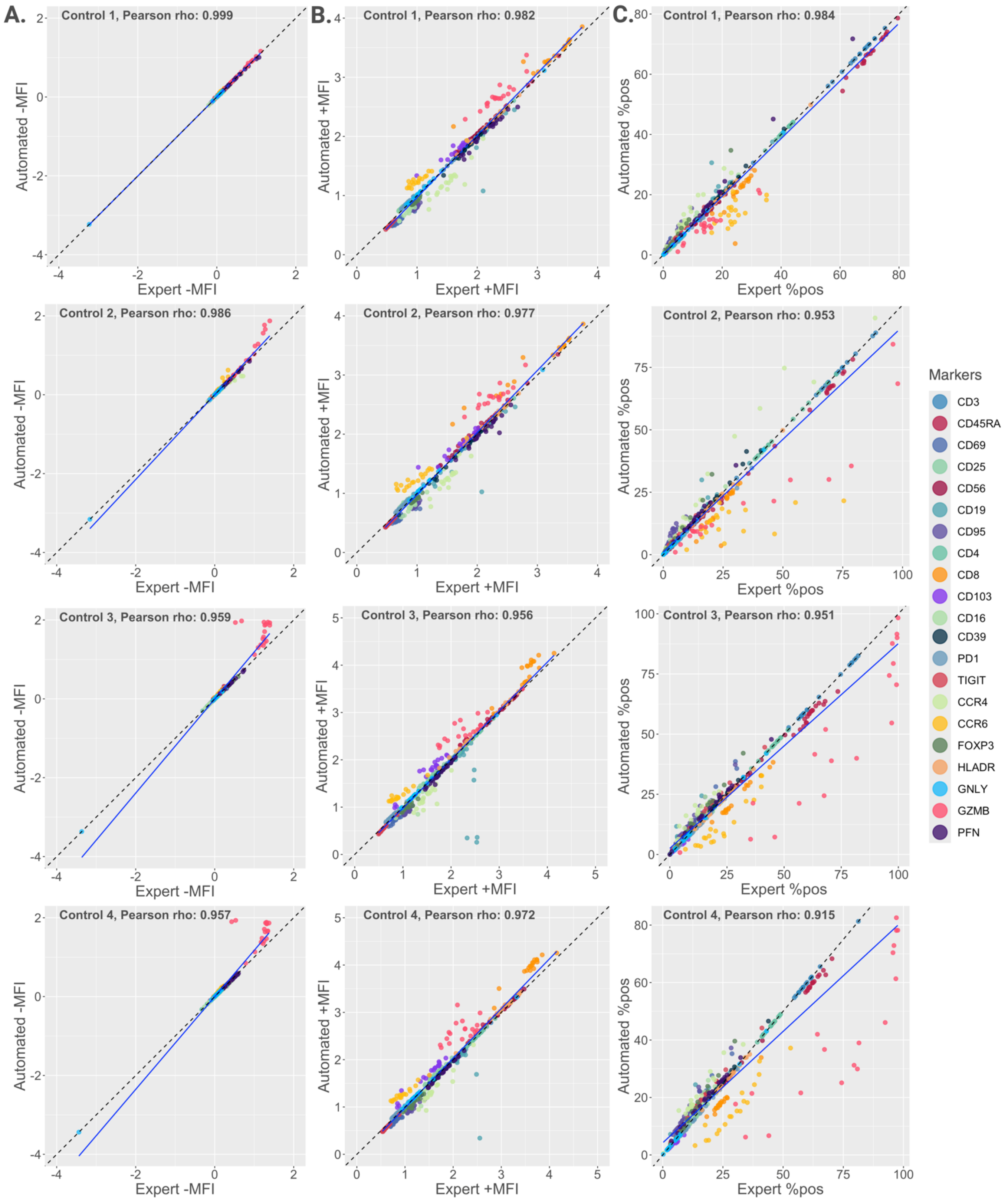


**Supplementary Figure 13. Comparison of IQR-based metrics in negative and positive populations separated based on automated and consensus expert thresholds (spectral dataset).** The automated and consensus expert thresholds were used to calculate the three metrics **(A)** -MFI, **(B)** +MFI, and **(C)** %pos for each marker and control sample. The resultant values are visualized using dot-plots comparing the consensus expert results (x-axis) with automated threshold results (y-axis). The dashed line indicates the identity line. Pearson's correlation coefficients (r) and their associated correlation lines are also represented.

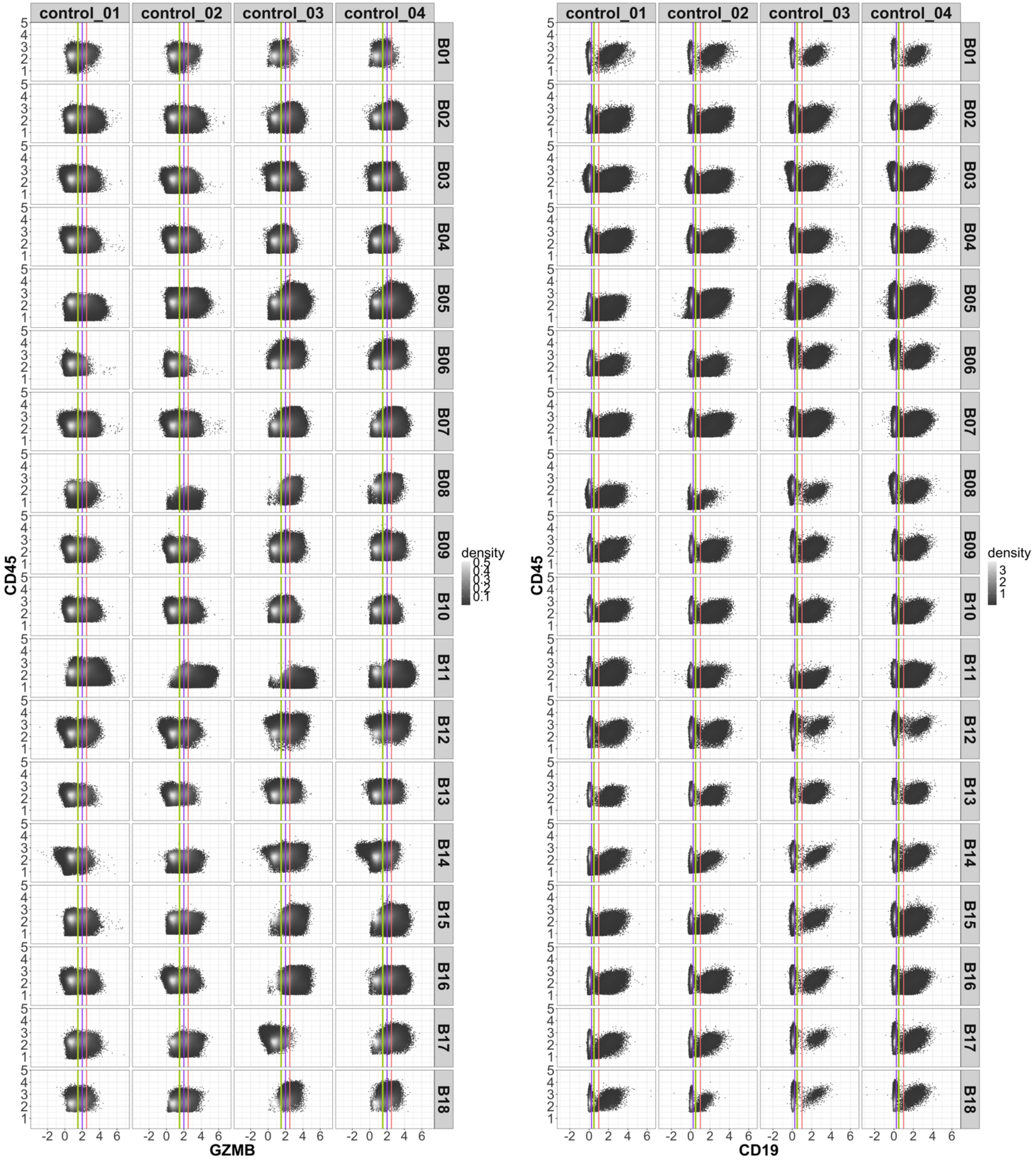


**Supplementary Figure 14.** Biaxial dot plots of GZMB (which showed variability between the %pos metric values for different expert and automated thresholds in Sup. Figs. 12, 13) and CD19 (which showed variable in +MFI) vs. CD45 across four control samples of the spectral dataset with minimum expert (green), consensus expert (yellow, behind green), maximum expert (coral) and automated (purple) threshold values denoted by the solid lines.

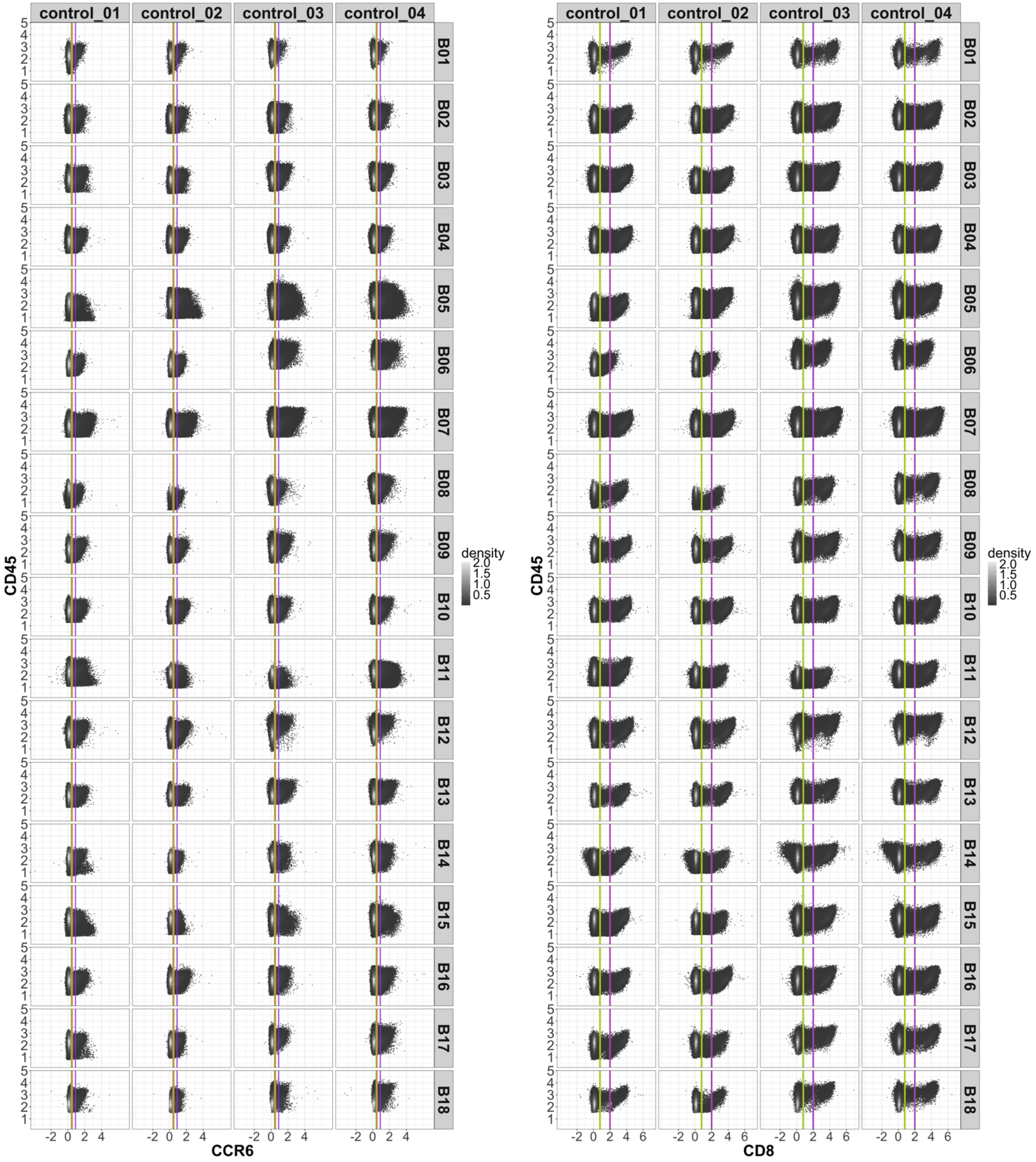


**Supplementary Figure 15.** Biaxial dot plots of CCR6 and CD8 vs. CD45 (which showed variability between the %pos metric values for different expert and automated thresholds in Sup. Figs. 12, 13) across four control samples of the spectral dataset with minimum expert (green), consensus expert (yellow, behind coral [left] and green [right]), maximum expert (coral, behind purple) and automated (purple) threshold values denoted by the solid lines.

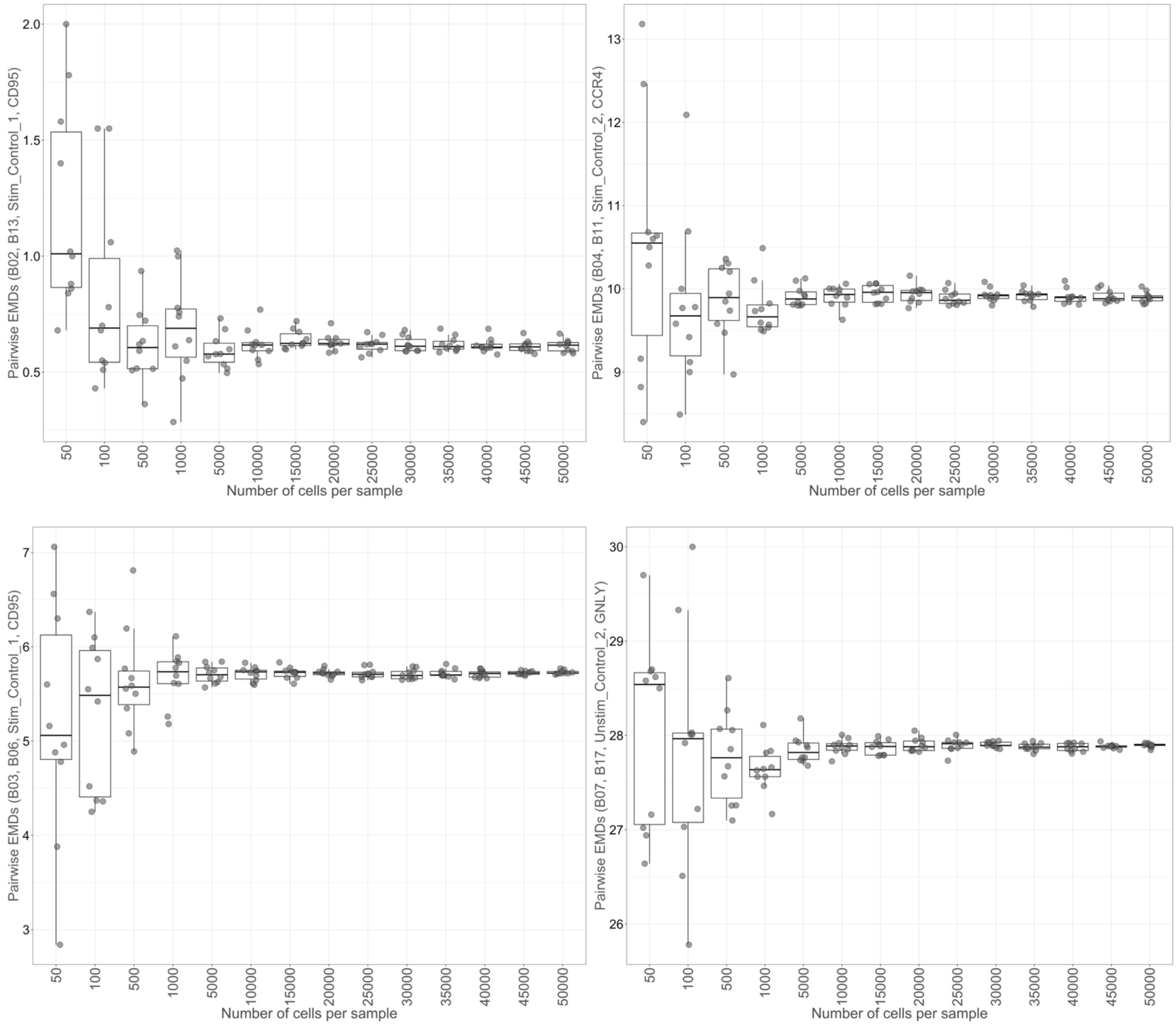


**Supplementary Figure 16. Evaluating the influence of number of randomly selected cells per sample on pairwise EMD measures in the spectral dataset.** Four different cases of pairwise EMDs for marker and control combinations of the spectral cytometry dataset. For each case, 50 to 50,000 cells per sample were randomly selected 10 times and EMDs calculated. EMD measures for each case are observed to stabilize anywhere between 10,000 to 20,000 cells per sample, dependent on the case evaluated.

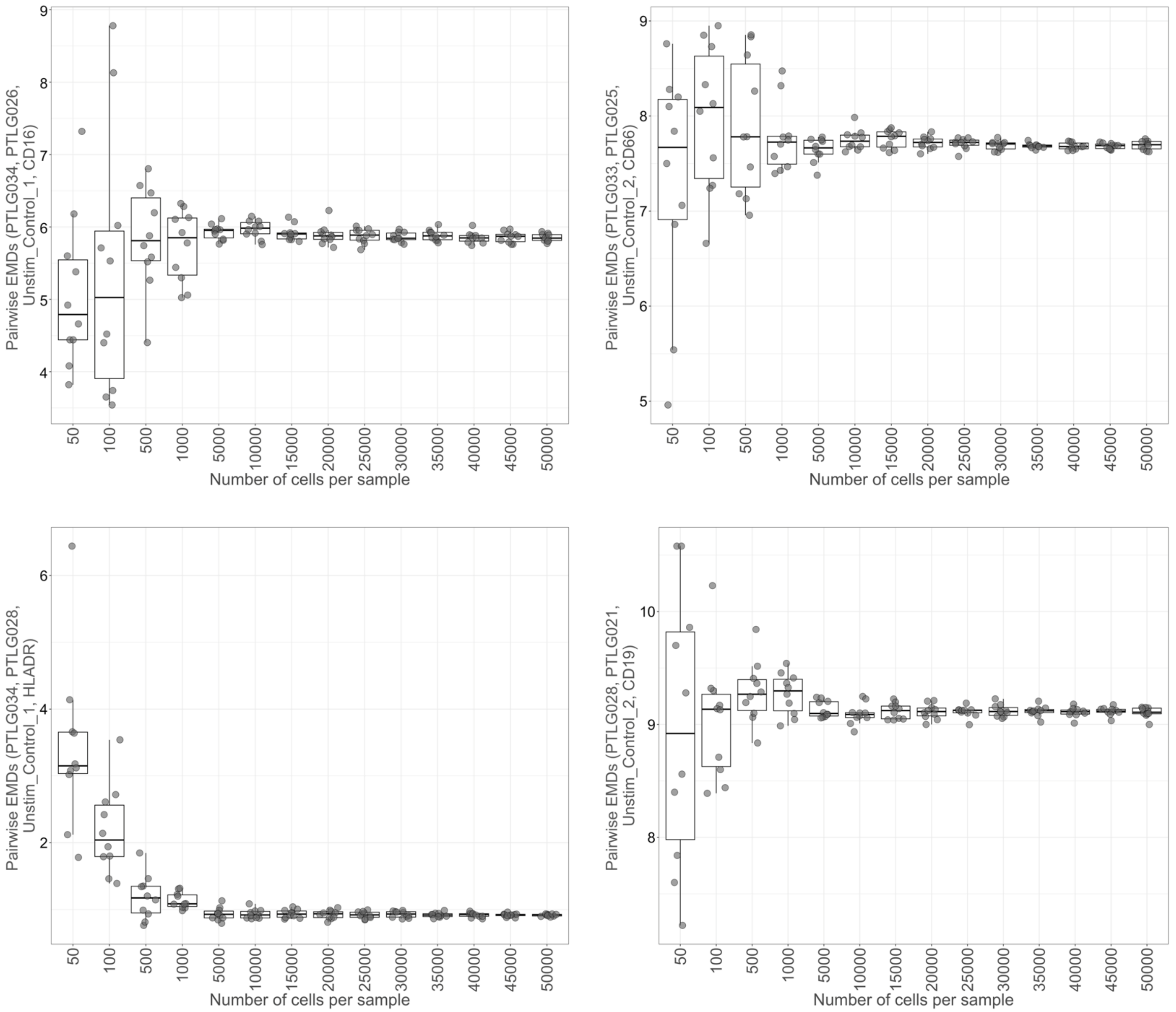


**Supplementary Figure 17. Evaluating the influence of number of randomly selected cells per sample on pairwise EMD measures in the CytoNorm dataset.** Four different cases of pairwise EMDs for marker and control combinations of the CytoNorm mass cytometry dataset. For each case, 50 to 50,000 cells per sample were randomly selected 10 times and EMDs calculated. EMD measures for each are observed to stabilize between 5,000 to 20,000 cells per sample (dependent on the case evaluated).

**Table S1.** Antibody panel used for the spectral dataset.

| Laser | P.Ch. | Dye | Antigen |
|---|---|---|---|
| **UV** | UV2 | BUV395-Tdm | CD4 |
| | UV7 | BUV496-Tdm | CD56 |
| | UV9 | BUV563-Tdm | CD3 |
| | UV11 | BUV661-Tdm | CD19 |
| | UV14 | BUV737-Tdm | CD95 |
| | UV16 | BUV805-Tdm | CD8 |
| **VIOLET** | V1 | BV421 | CD103 |
| | V2 | SB436 | |
| **405** | V3 | P. Blue | **FOXP3*** |
| | V4 | BV480 | CD16 |
| | V7 | BV510 | CD14 |
| | V8 | BV570 | |
| | V10 | BV605-Tdm | CD69 |
| | V11 | BV650-Tdm | CCR6 |
| | V13 | BV711-Tdm | **Perforin*** |
| | V14 | BV750-Tdm | HLA-DR |
| | V15 | BV785-Tdm | |
| **BLUE** | B1 | BB515 | |
| **488** | B2 | Alexa488 | CCR7 |
| | B3 | SB550 | CD45 |
| | B5 | PerCP | |
| | B8 | PerCP-Cy5.5 | |
| | B10 | PerCP/eF710 | **TIGIT** |
| **YELLOW** | YG1 | PE | CD25 |
| **561** | YG3 | PE/Dzle594 | CD39 |
| | YG5 | PE/Cy5 | CD45RA |
| | YG7 | PE/Cy5.5 | |
| | YG9 | PE/Cy7 | PD1 |
| **RED** | R1 | APC | CCR4 |
| **640** | R2 | Alexa647 | **GNLY*** |
| | R4 | Alexa700 | **HELIOS*** |
| | R7 | APC/Fire750 | **GZMB*** |

* denotes intracellular staining.
